# Capturing dynamic phage-pathogen coevolution by clinical surveillance

**DOI:** 10.1101/2025.01.29.635557

**Authors:** Yamini Mathur, Caroline M. Boyd, Jeannette E. Farnham, Md Mamun Monir, Mohammad Tarequl Islam, Marzia Sultana, Tahmeed Ahmed, Munirul Alam, Kimberley D. Seed

## Abstract

Bacteria harness diverse defense systems that protect against phage predation^1^, many of which are encoded on horizontally transmitted mobile genetic elements (MGEs)^2^. In turn, phages evolve counter-defenses^3^, driving a dynamic arms race that remains underexplored in human disease contexts. For the diarrheal pathogen *Vibrio cholerae*, a higher burden of its lytic phage, ICP1, in patient stool correlates with reduced disease severity^4^. However, direct molecular evidence of phage-driven selection of epidemic *V. cholerae* has not been demonstrated. Here, through clinical surveillance in cholera-endemic Bangladesh, we capture the acquisition of a parasitic anti-phage MGE, PLE11, that initiated a selective sweep coinciding with the largest cholera outbreak in recent records. PLE11 exhibited potent anti-phage activity against co-circulating ICP1, explaining its rapid and dominating emergence. We identify PLE11-encoded Rta as the novel defense responsible and provide evidence that Rta restricts phage tail assembly. Using experimental evolution, we predict phage counteradaptations against PLE11 and document the eventual emergence and selection of ICP1 that achieves a convergent evolutionary outcome. By probing how PLEs hijack phage structural proteins to drive their horizontal transmission while simultaneously restricting phage tail assembly, we discover that PLEs manipulate tail assembly to construct chimeric tails comprised of MGE and phage-encoded proteins. Collectively, our findings reveal the molecular basis of the natural selection of a globally significant pathogen and its virus in a clinically relevant context.

## Introduction

The infectious diarrheal disease cholera, caused by the pathogen *Vibrio cholerae*, remains a significant threat to global public health. The Bay of Bengal is considered the source of the ongoing seventh cholera pandemic, where seventh pandemic El Tor (7PET) sublineages evolve and spread to vulnerable nations across the globe^5^. Unraveling the factors influencing the evolution and selection of 7PET strains is challenging, as they result from the complex interplay of genetic changes, particularly the flux of novel mobile genetic elements (MGEs), and selective advantages that can be difficult to ascertain. Recent metagenomic analyses in cholera-endemic Bangladesh indicate that higher ratios of ICP1, the predominant lytic phage preying on *V. cholerae* in the context of disease^6^, correlate with reduced risk of severe disease in patients^4^. This suggests that the acquisition of phage resistance could contribute to outbreak severity and influence the evolution of pandemic lineages. However, direct molecular evidence of this link is lacking, as are mechanistic insights into how the dynamic oscillations of phage susceptibility and resistance unfold in nature.

Among the fluctuating MGEs found in 7PET *V. cholerae* are a family of phage satellites called phage-inducible chromosomal island-like elements (PLEs) that play a crucial role in defending against ICP1 predation^7^. PLEs have evolved intricate defense mechanisms that disrupt the phage lifecycle while exploiting phage machinery and structural components to package themselves into modified viral particles, thereby blocking phage transmission^7–10^. PLEs are highly specialized in safeguarding *V. cholerae* populations from predation by ICP1, but phage counteradaptations can neutralize their potent anti-phage activity. Analyses of sparsely collected ICP1 isolates have revealed three anti-PLE mechanisms the phage uses to overcome PLE-mediated hijacking and restore phage propagation. These mechanisms vary among phage isolates^6^ and mediate nucleolytic degradation of the PLE genome through distinct mechanisms (Fig. 1a). The phage-encoded origin-directed nuclease (Odn)^11^ and attachment-directed inhibitor (Adi)^12^ antagonize specific subsets of the ten PLE variants discovered to date^13^, targeting sequence variants of the PLE origin of replication and integrase, respectively. A remarkable adaptation in ICP1 is its co-option of CRISPR-Cas, which replaces *odn* in the same genomic locus and provides broad-spectrum counter-defense against all PLEs tested to date^11,14^. Interestingly, genomic analyses have documented the temporal flux of distinct PLE variants in epidemic *V. cholerae*^13^. However, without knowledge of co-circulating ICP1 genotypes, the molecular factors driving such epidemiological patterns remain unclear. Building on the foundational understanding of PLE-ICP1 conflict, we harnessed high-resolution clinical surveillance in Bangladesh to capture and understand the emergence and selection of epidemic *V. cholerae* and ICP1 in a region critical to the dynamics of pandemic cholera.

**Figure 1.**
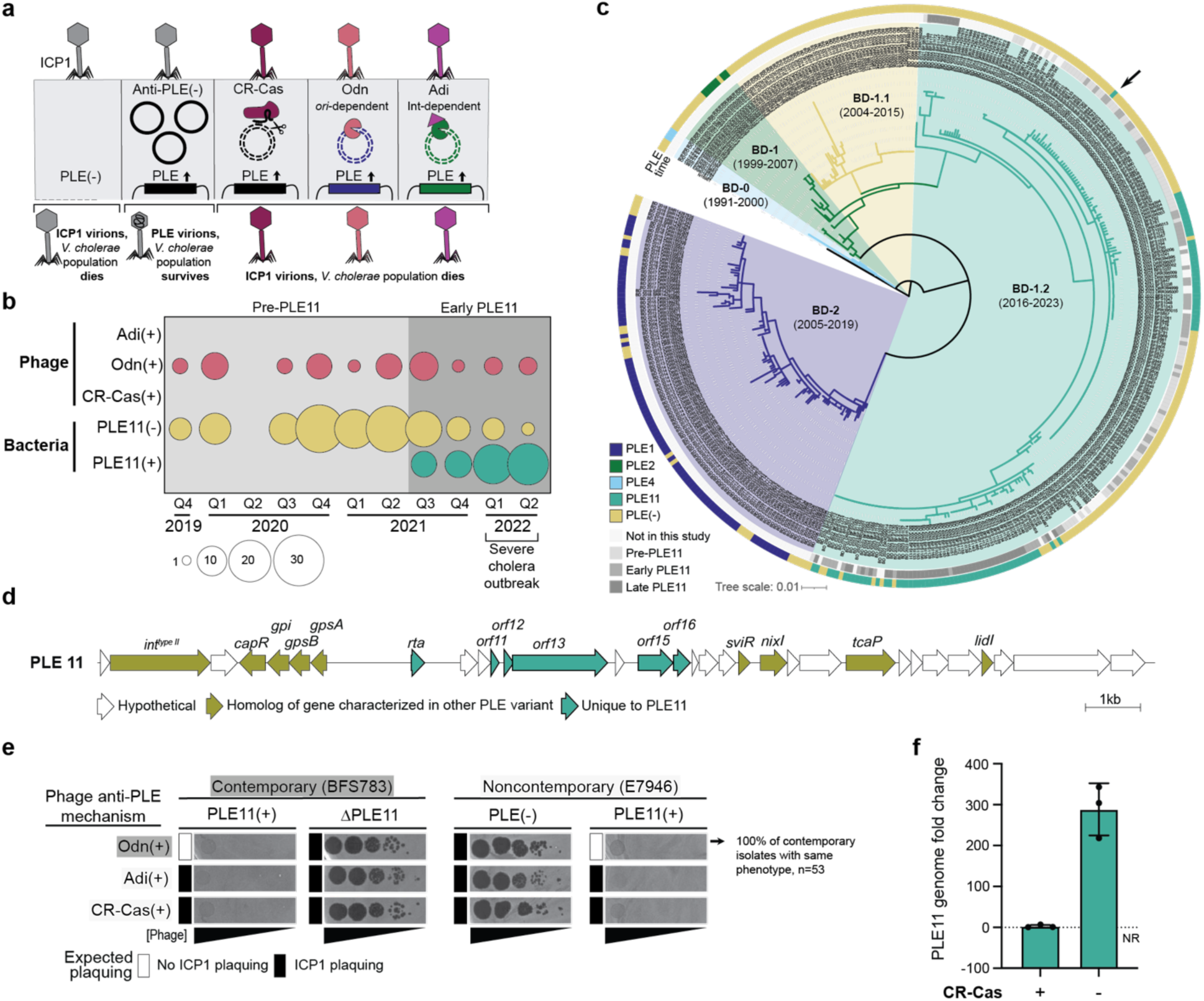
Clinical surveillance reveals the acquisition and selection of PLE11 in *V. cholerae* that restricts ICP1 despite anti-PLE mechanisms. **a)** Summary model of validated infection outcomes of PLE(+) *V. cholerae* with ICP1 isolates encoding anti-PLE mechanisms. Upon ICP1 infection, PLE excises from the chromosome, replicates, and hijacks ICP1-encoded structural proteins to package its genome for horizontal transmission while producing inhibitors that synergistically abolish phage production^8–10^. ICP1 has evolved three distinct anti-PLE mechanisms^11,12,14^ that destroy the PLE genome, restoring phage production: CRISPR-Cas (CR-Cas) cleaves diverse PLEs, while the origin-directed nuclease (Odn) and the attachment-directed inhibitor (Adi) antagonize a subset of PLE variants based on the sequence of the origin of replication or the presence of a Type II integrase (Int), respectively. **b)** Sampling of 191 *V. cholerae* (n=143 whole genome sequenced) and 53 ICP1 isolates (n=17 whole genome sequenced) from cholera patient stool samples in Bangladesh depicting isolate counts by genotype. Q1 = January-March, Q2 = April-June, Q3 = July-September, Q4 = October-December. The light gray shading indicates the period before PLE11 (pre-PLE11), and the medium gray shading indicates the first year after the initial detection of PLE11 (early PLE11 period). **c)** Maximum likelihood phylogeny of 431 assembled *V. cholerae* strains from Bangladesh using reference strain N1961 as an outgroup. Lineages, as in Monir et al.^15^, are highlighted with background colors, and the isolation time range for isolates in each lineage is indicated. Colored rings outside the tree indicate the time of the isolate’s isolation if sequenced as part of this study, which matches the time shading from panel b (innermost ring), and the presence and identity of a PLE are indicated by the legend (outermost ring). The first clinical isolate harboring PLE11 in our surveillance, BFS783, used in panel e, is indicated by an arrow. The scale bar represents the number of nucleotide substitutions per site. **d)** Genetic organization of PLE11 drawn to scale. Arrows represent genes colored according to the legend. **e)** Plaquing of tenfold serially diluted ICP1 phage isolates, with the anti-PLE mechanism indicated, on lawns of *V. cholerae*. The gray background is the bacterial lawn, and the dark spots are zones of killing. The expected plaquing phenotype is indicated for each host-phage pair. The plaquing phenotype for Odn(*+*) phage on PLE11(+) E7946 is representative of all ICP1 isolates recovered during the pre-PLE11 and early PLE11 periods (n=53) (Extended Data Fig. 2a, Supplementary Fig. 1). CRISPR-Cas(+) and Adi(+) ICP1 isolates are historically collected isolates. **f)** Replication of PLE11 in *V. cholerae* calculated as the fold change in PLE11 DNA copy by qPCR 20 minutes post-infection relative to immediately pre-infection by otherwise isogenic CRISPR-Cas(+/-) ICP1. The bar represents the mean; each dot represents a biological replicate; the error bars indicate the standard deviation, and the dotted line at a fold change of 1 indicates no replication (NR).

### PLE11 fuels selective sweep of *V. cholerae*

Between October 2019 and June 2022, we collected and analyzed 516 stool samples from patients with suspected cholera from the megacity of Dhaka (n = 418) and the small coastal village of Mathbaria (n=98) in Bangladesh. Clinical *V. cholerae* and phage isolates were subjected to phenotypic analysis and genome sequencing to ascertain emerging patterns in their coevolutionary arms race (Supplementary Tables 1 and 2). Notably, this surveillance period coincided with a massive cholera outbreak in March-April 2022, during which the icddr,b Dhaka hospital treated over 42,00 patients (Fig. 1b)^15^. We sequenced the genomes of 143 *V. cholerae* isolates from unique stool samples during this period as well as 53 from the following 15 months (discussed below) and integrated them with 235 previously published genomes of Bangladeshi *V. cholerae* O1 isolates collected since 1991. Core-genome phylogenetic analysis of the combined genome set (n=431) revealed that strains from this surveillance period belonged to the BD-1.2 sublineage of 7PET wave-3 strains (Fig. 1c). This lineage displaced the previously locally dominant PLE1(+) BD-2 lineage in 2018 and was initially comprised of PLE(-) strains^13,15^.

In September 2021, we captured the acquisition of a novel variant of the MGE PLE, designated PLE11, in the BD-1.2 lineage (Figs. 1b and 1d). Compellingly, PLE11 was acquired in *V. cholerae* amid a backdrop of co-circulating ICP1 isolates invariably encoding Odn as their sole anti-PLE mechanism; further, within 9 months of its initial detection, PLE11 was present in 91% of *V. cholerae* isolates (Fig. 1b). PLE11 does not possess the origin of replication that is sensitive to ICP1’s anti-PLE nuclease Odn (Extended Data Fig. 1a), indicating this MGE would likely protect *V. cholerae* from co-circulating ICP1 via conserved and previously established PLE-encoded defenses against ICP1 (Gpi, TcaP, NixI)^8–10^ (Fig. 1d and Extended Data Fig. 1a). To experimentally evaluate whether PLE11 can protect *V. cholerae* from Odn(+) ICP1 infection, we first used the earliest PLE11(+) clinical isolate detected in our surveillance, BFS783. We constructed a deletion of the PLE in BFS783 and, in parallel, introduced PLE11 into a noncontemporary, phage-sensitive, and PLE(-) *V. cholerae* strain, E7946. We then assessed PLE11-mediated inhibition of contemporary ICP1 isolates (n=53), all of which were Odn(+) and recovered only from stool samples with PLE(-) *V. cholerae*. As anticipated, PLE11 was both necessary and sufficient to abolish plaque formation for all co-circulating phage isolates (Fig. 1e, Extended Data Figs. 1b and 2, Supplementary Fig. 1). In comparison, the closely related PLE(-) clinical isolate BFS948 was susceptible to all Odn(+) co-circulating phages (Extended Data Fig. 2b). As such, during the severe cholera outbreak in 2022, no ICP1 isolates were detected that were capable of preying on the dominant PLE11(+) strains that accounted for the vast majority of the circulating *V. cholerae*. These data demonstrate that the novel MGE variant PLE11 conferred a selective advantage to *V. cholerae in situ* and provide direct evidence that phage resistance contributes to the succession of epidemic strains.

Given the lack of co-circulating phages with an effective means to combat PLE11 during this period, we turned to our historical collection of ICP1 isolates to evaluate predicted infection outcomes based on the known specificity of alternative anti-PLE mechanisms Adi and CRISPR-Cas. We hypothesized that PLE11 would be susceptible to CRISPR-Cas(+) or Adi(+) phages, given the universal efficacy of CRISPR-Cas against all previously characterized PLE variants^7,13^ and its Type II integrase. Surprisingly, PLE11 protected *V. cholerae* even when infected by phages encoding CRISPR-Cas or Adi (Fig. 1e, Extended Data Fig. 1b). We omitted the possibility of a PLE11-encoded anti-CRISPR because the PLE11 genome was still degraded upon CRISPR-Cas(+) phage infection (Fig. 1f). This indicated that, unlike all previously characterized PLE variants, PLE11 employs a novel anti-ICP1 defense mechanism that functions despite nucleolytic targeting of the PLE genome. Together, our results show the rapid and dominating emergence of PLE11 that not only protected *V. cholerae* from infection by co-circulating ICP1 but also maintained phage defense in the face of all previously characterized anti-PLE mechanisms.

### Predicting ICP1’s counteradaptations to PLE11

For ICP1 to sustain its long-term association with epidemic *V. cholerae*, we expected ICP1 to eventually evolve to circumvent defense by PLE11. To predict such an adaptation in nature, we began by investigating the molecular basis of PLE11’s unique capacity to restrict ICP1 despite its anti-PLE mechanisms. To this end, we focused on genes unique to PLE11 (Fig. 1d). Deletion of one such gene, *rta,* rendered PLE11(+) *V. cholerae* susceptible to ICP1 infection (Fig. 2a, Extended Data Fig. 3a). Importantly, ICP1 still required an appropriate anti-PLE mechanism to productively infect Δ*rta* PLE11 *V. cholerae*, consistent with PLE11 encoding known defenses against ICP1^8–10^ and PLE11’s predicted susceptibility to CRISPR-Cas and Adi (Fig. 2a, Extended Data Fig. 3a). Furthermore, ectopic expression of *rta* alone was sufficient to restrict all 53 contemporary ICP1 isolates to a comparable degree as PLE11 (Extended Data Fig. 2a, Supplementary Fig. 1).

**Figure 2.**
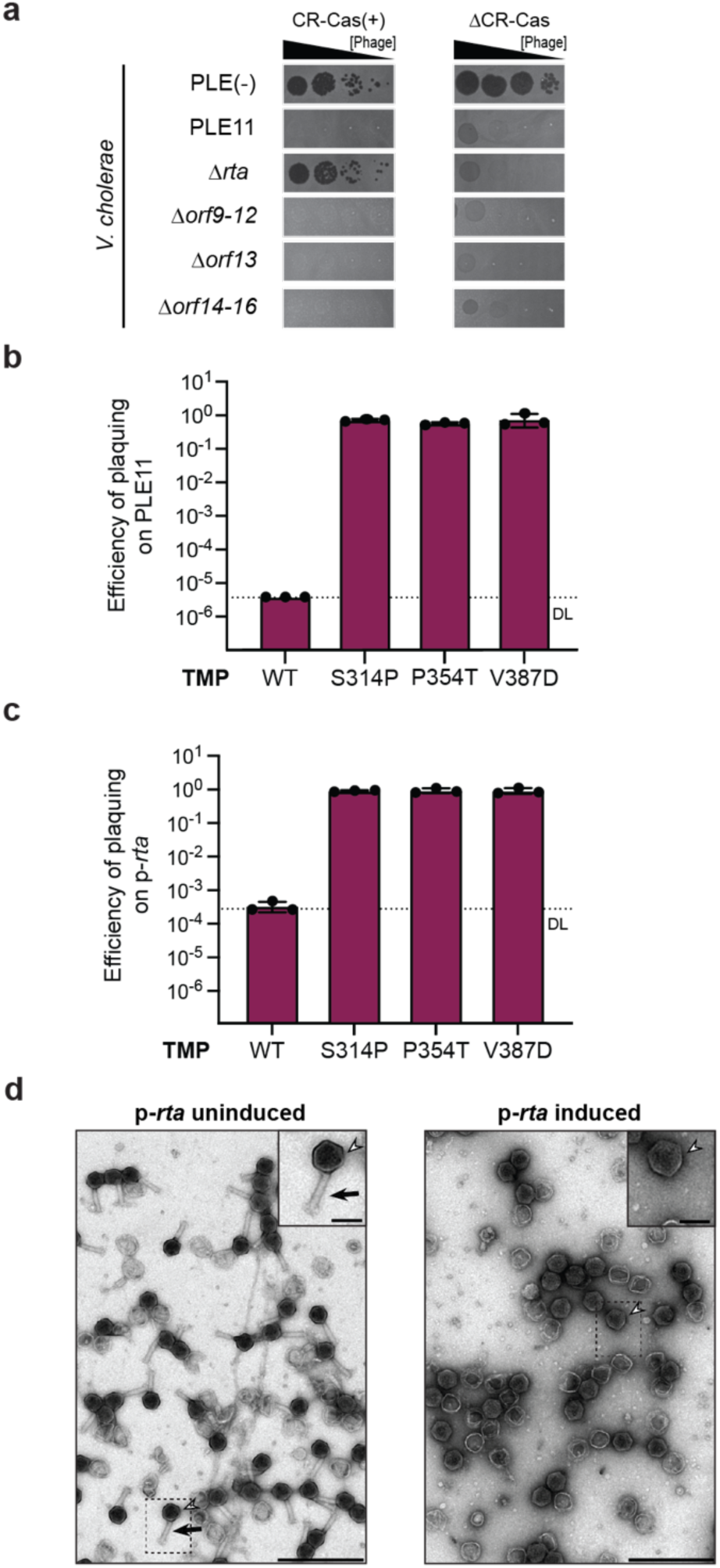
Experimental evolution of ICP1 resistant to PLE11 reveals novel tail targeting defense protein, Rta, and predicts counteradaptations needed for ICP1 to persist in nature. **a)** Plaquing of tenfold serially diluted ICP1 CRISPR-Cas(+/-) on lawns of *V. cholerae* strain E7946 and its PLE11(+) and PLE11 mutant derivatives. **b** and **c)** Efficiency of plaquing of wild type (WT) CRISPR-Cas(+) ICP1 and experimentally evolved derivatives isolated on PLE11(+) *V. cholerae*, indicated by the relevant substitution in the tape measure protein (TMP), on (b) PLE11(+) *V. cholerae* relative to a permissive PLE(-) strain or (c) on Rta-expressing *V. cholerae* relative to permissive *V. cholerae* harboring an empty vector. The bar represents the mean; each dot represents a biological replicate; the error bars indicate the standard deviation, and DL indicates the detection limit. **d)** Representative transmission electron micrographs (TEMs) of particles produced following ICP1 infection of PLE(-) *V. cholerae* with *rta* expressed from a low copy plasmid (+/-) inducer. The arrowheads indicate DNA-filled capsids, and the arrow indicates a virion tail. The scale bars are 500nm and 100nm for the zoomed-out views and insets, respectively.

Rta is a small 80 amino acid protein for which we could not identify homologs nor ascertain a potential function using primary sequence or predicted structure. To gain insight into Rta’s mechanism of phage restriction in its native genomic context, we evolved CRISPR-Cas(+) ICP1 mutants that escaped PLE11. All escape mutants had single nonsynonymous mutations in *gp79* or *gp81* (Fig. 2b and Supplementary Table 3), the transcripts of which are spliced to produce ICP1’s tape measure protein (TMP)^16^. These phages were also insensitive to ectopic expression of Rta (Fig. 2c), indicating TMP substitutions are sufficient to escape Rta when expressed natively or synthetically. Strengthening these findings, we repeated the experimental evolution approach using a phylogenetically distinct Adi(+) ICP1 isolate and obtained parallel results in which all escape mutants harbored substitutions within the TMP (Supplementary Table 4).

In contractile phages like ICP1, the TMP occupies the core of the phage tail and serves as the essential assembly scaffold determining tail length^17^. To substantiate the hypothesis that Rta targets ICP1’s TMP, we examined the morphology of ICP1 virions produced from infection of *V. cholerae* ectopically expressing Rta by transmission electron microscopy (TEM). With the induction of Rta, we observed an abundance of genome-filled capsids lacking tails (Fig. 2d, Extended Data Fig. 3b). Together, these results identify Rta as a novel phage defense protein and suggest that Rta disrupts ICP1 infection by targeting its TMP, resulting in defective tailless virions incapable of propagation. Most notably, Rta uniquely enables PLE11 to protect *V. cholerae* populations despite the nucleolytic degradation of the PLE genome mediated by ICP1’s anti-PLE mechanisms. Collectively, these data provide a predicted set of adaptations needed for ICP1 to circumvent a recently acquired anti-phage MGE during a natural outbreak of *V. cholerae*.

### Capturing ICP1’s counteradaptations in nature

Our laboratory evolution studies revealed that ICP1 could evade PLE11-mediated defense through a combination of a nucleolytic anti-PLE mechanism (CRISPR-Cas or Adi) to curb all conserved PLE-encoded defenses and a substitution in the tape measure protein (TMP) to circumvent Rta activity (Fig. 2). Our continued surveillance offered a rare opportunity to evaluate how the genetic basis of adaptation might differ between experimental and clinical phage populations. From July 2022 to September 2023, we collected 189 stool samples to monitor the persistence of PLE11 in *V. cholerae* and track ICP1’s evolutionary response to this novel mobile phage defense element (Supplementary Tables 1 and 2). PLE11 was detected in 100% (49/49) of BD-1.2 sublineage isolates in 2023, rising from 20% (17/84) in 2021 (Fig. 3a). PLE11(+) strains correspond to two distinct phylogenetic clusters within BD-1.2, suggesting at least two independent acquisitions of this anti-phage MGE (Fig. 1c). The complete replacement of PLE(-) strains with PLE11(+) *V. cholerae* is consistent with the profound fitness advantage conferred by this novel mobile phage defense element. By 2023, this genotype shift in *V. cholerae* also coincided with the disappearance of Odn(*+*) ICP1 phages from stool samples (Fig. 3a). As we observed from the pre-PLE11 and early-PLE11 periods, no Odn(+) isolates from the late PLE11 period were recovered from patient stool samples in which we isolated PLE(+) *V. cholerae*. These Odn(+) isolates also failed to propagate on PLE11(+) *V. cholerae* and were restricted by Rta (Fig. 3b and c, Extended Data Fig. 5 and Supplementary Fig. 2). Significantly, 11 months after PLE11 was initially detected, ICP1 isolates capable of propagating on PLE11(+) *V. cholerae* emerged (Fig 3a). These phages also exhibited resistance to Rta-mediated restriction (Fig. 3b, Extended Data Fig. 5 Supplementary Fig. 2.), demonstrating that ICP1 coevolved in nature to circumvent phage defense mediated by the novel PLE variant and, specifically, Rta activity.

**Figure 3.**
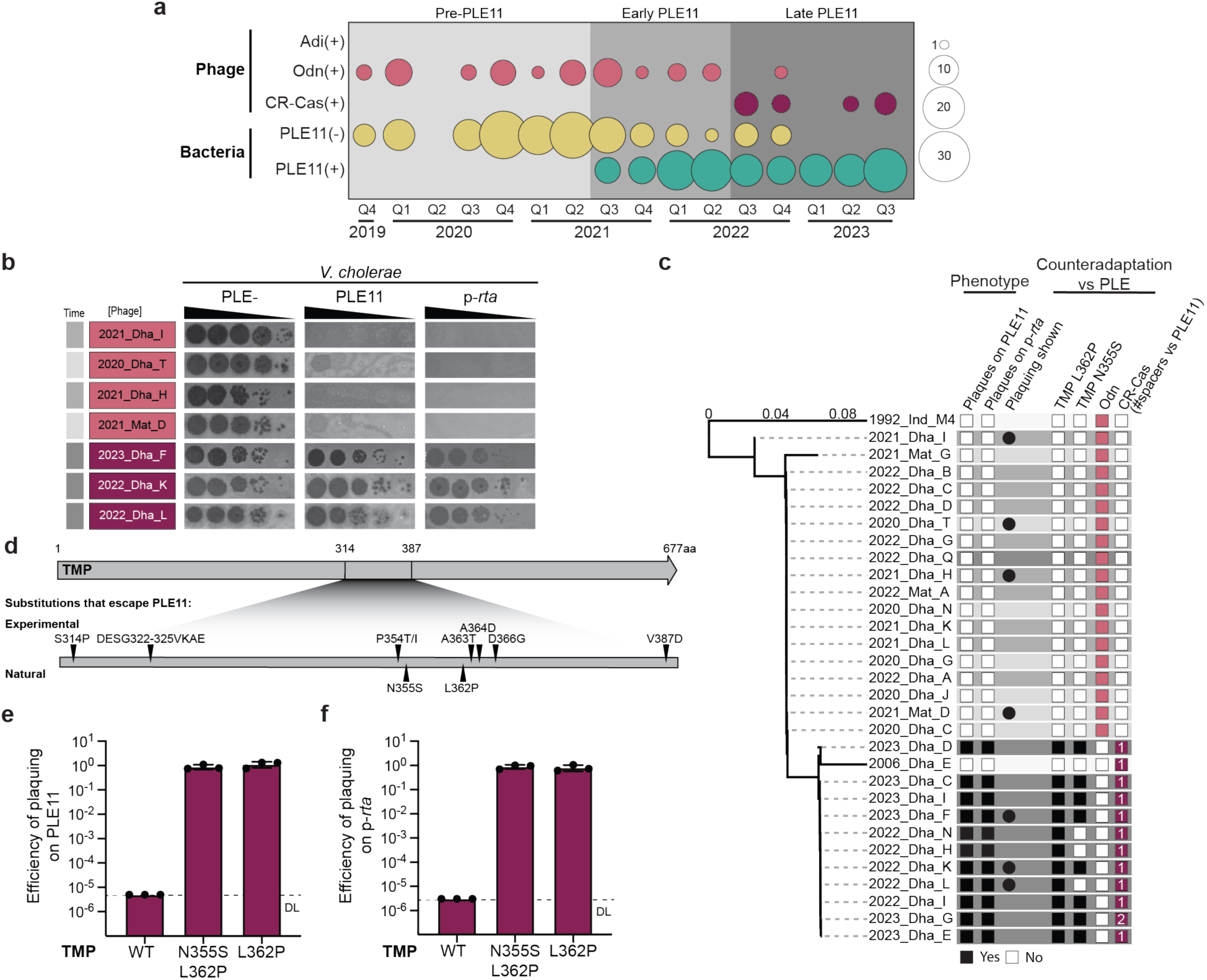
Continued surveillance documents the eventual emergence of ICP1 with counteradaptations against PLE11 and Rta. **a)** Sampling of 275 *V. cholerae* (n=196 whole genome sequenced) and 75 ICP1 (n=29 whole genome sequenced) from cholera patient stool samples in Bangladesh depicting isolate counts by genotype. The light gray shading indicates the period before PLE11 (pre-PLE11), the medium gray shading indicates the first year after the initial detection of PLE11 (early PLE11 period), and the dark gray indicates the time period with CRISPR-Cas(+) ICP1 co-circulating with PLE11(+) *V. cholerae* (late PLE11 period). **b)** Representative plaquing of tenfold serially diluted ICP1 phage isolates with the anti-PLE mechanism indicated (Odn (pink) or CR-Cas (maroon)) on lawns of *V. cholerae* strain E7946, E7946 PLE11(+) and E7946 with a low-copy plasmid expressing PLE11 Rta (p-*rta*). **c)** Phylogeny of whole genome sequenced ICP1 phages (29 from this study with previously identified isolates 1992_Ind_M4 and 2006_Dha_E included as references) built by comparing their translated open reading frames (using the tBLASTx algorithm from ViPTree). Standardized ICP1 names are shown (which include the year and location of isolation: Dha = Dhaka Bangladesh, Mat = Mathbaria Bangladesh, Ind = India). Gray shading corresponds to the time of the phage’s isolation if sequenced as part of this study, which matches the time shading from panel a. Empty boxes represent no phenotype or absent genotype, while filled boxes represent yes phenotype or present anti-PLE mechanism (Odn (pink) or CRISPR-Cas (maroon)). For CRISPR-Cas, the number of spacers that are >96% identical to PLE11 are reported within the colored box. Note: 2023_Dha_G carries a novel spacer uniquely targeting PLE11. **d)** Schematic of the region of ICP1’s tape measure protein (TMP) with indicated amino acid substitutions from experimentally evolved phages selected on PLE11(+) *V. cholerae* and from CRISPR-Cas(+) ICP1 isolates co-circulating with PLE11 (+) *V. cholerae* in clinical samples. **e** and **f)** Efficiency of plaquing of wild type (WT) CR-Cas(+) ICP1 and lab-engineered derivatives that encode one or both substitutions seen in the TMP of natural clinical phage isolates that co-circulate with PLE11 on (e) PLE11(+) *V. cholerae* relative to a permissive PLE(-) strain or (f) on Rta-expressing *V. cholerae* relative to permissive *V. cholerae* harboring an empty vector. The bar represents the mean; each dot represents a biological replicate; the error bars indicate the standard deviation, and the DL indicates the detection limit.

To identify the genetic basis of these adaptations, we examined the anti-PLE mechanism(s) in these ICP1 isolates and found they exclusively encoded CRISPR-Cas, with each phage encoding at least one spacer targeting the PLE11 genome (Figure 3a and c). This genotype shift was not the result of exchanging Odn for CRISPR-Cas in the ICP1 population but instead reflected a replacement of the predominantly clonal Odn(+) ICP1 population with phylogenetically distinct, largely clonal CRISPR-Cas(+) ICP1 isolates (Fig. 3c and Extended Data Fig. 4). Consistent with observations from experimentally evolved phages, we found that CRISPR-Cas(+) ICP1 isolates co-circulating with PLE11 harbored one or more amino acid substitutions in the TMP. Mapping the substitutions in evolved phage from clinical specimens revealed they localized within the same 73 amino acid region of the TMP as identified in our experiments (Fig. 3d). These substitutions, L362P (in all isolates) and N355S (in 79%), were adjacent to residues that conferred escape from PLE11 and ectopically expressed Rta in laboratory conditions, though they were distinct from those arising in experimental conditions (Fig. 3d, Extended Data Fig 5, and Supplementary Fig. 2). To confirm the functional role of naturally evolved TMP substitutions, we engineered L362P or L362P + N355S into the CRISPR-Cas(+) Rta-sensitive historical isolate of ICP1 used in experimental evolution. Both variants were sufficient to evade phage defense mediated by PLE11 via Rta (Fig. 3e and f), demonstrating that ICP1 evolved in nature under selection pressures imposed by a novel phage defense in *V. cholerae*.

In summary, our surveillance revealed the predicted shift from phages with the ineffective anti-PLE mechanism Odn to phages with CRISPR-Cas that overcomes PLE11’s conserved anti-phage defenses and substitutions in the TMP to counter PLE11’s unique Rta-mediated phage defense. While experimental evolution studies could not select for CRISPR-Cas(+) isolates from an Odn(+) population— due to the lack of pre-existing variation in genetically homogenous stocks—natural populations likely maintain some level of genetic diversity, enabling selection of rare alleles and oscillations of the dominant genotypes. Together, our results document the coevolution of a lytic phage in a clinically relevant context to counter a newly acquired phage defense and demonstrate a convergent evolutionary outcome in nature that mirrors experimental studies.

### PLEs evade Rta via chimeric tail assembly

All characterized dsDNA phage satellites hijack phage tails for virion assembly. Thus, our data indicating that Rta interferes with phage tail assembly (Fig. 2) is seemingly at odds with the current paradigm of how satellites restrict phages while promoting their horizontal transmission using stolen structural components. The most extensively characterized PLE variant, PLE1, transduces in virions with contractile tails^18^, dependent on the ICP1 receptor^7^, consistent with hijacking ICP1-encoded tails. Similarly, TEM of purified PLE11 virions revealed particles with modified capsids, attributed to the PLE-encoded capsid scaffold protein TcaP^9^, and contractile tails (Fig. 4a). We inferred from these observations that PLE11 must have a mechanism to assemble tailed virions in the presence of Rta.

**Figure 4.**
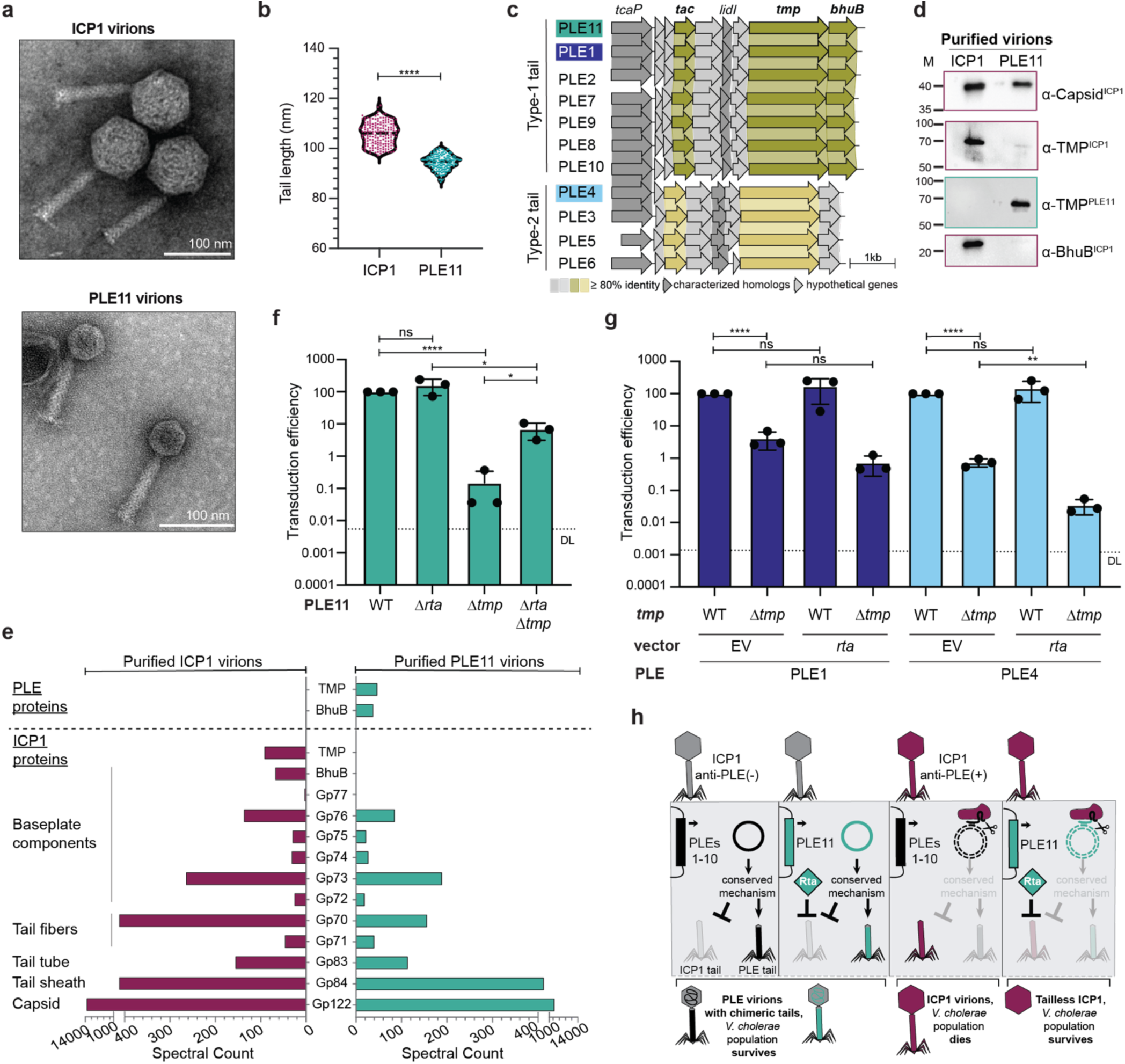
Novel incorporation of satellite components into chimeric tails by PLEs allows Rta to restrict ICP1’s tail assembly. **a)** Representative transmission electron micrographs (TEM) of ICP1 and PLE11 virions. The scale bar = 100 nm. **b)** Lengths of ICP1 and PLE11 virion tails analyzed from TEM images of purified virions. Empirically measured ICP1 tail lengths of 105.9 ± 0.35 nm and PLE tail lengths of 94.48 ± 0.22 nm) (Mean ± SEM) match the predicted ranges based on protein length (ICP1’s 677 amino acid TMP, PLE’s 582 amino acid TMP) and the reported 1.6 Å per amino acid residue^19^. Each dot represents the tail length of an individual particle (n=170 for each ICP1 and PLE), with the mean shown as a horizontal line; statistical significance was assessed using Student’s t-test (****p <0.0001). **c)** Bioinformatic analysis of *V. cholerae* PLEs’ open reading frames (arrows) using tBLASTX. Similar gene products found in multiple PLEs have links drawn between them and are shaded if >80% identity. Previously characterized PLE1 proteins and homologs in other PLEs are indicated. Homologs of predicted tail proteins, including the tail assembly chaperone (TAC), tape measure protein (TMP), and baseplate hub (BhuB), are categorized into groups based on sequence identity. Homologs exhibit <25% identity between groups, while within each group, they share >90% identity. All PLE-encoded tail assembly-related proteins (TAC, TMP, and BhuB) have 19-41% identity to the equivalent counterparts in ICP1 (Supplementary Table 10). **d)** Western blot analysis of purified ICP1 and PLE11 virions for indicated tail components and ICP1’s major capsid protein (known to comprise the capsids of PLE and ICP1 virions^9^). **e)** Total protein from purified ICP1 and PLE11 virions analyzed for structural composition using collisional-induced dissociation mass spectrometry. The spectral counts (x-axes) correspond to a subset of proteins (y-axis) from the PLE11 (top) and ICP1 proteomes (bottom) from ICP1 (left) and PLE11 (right) virions, respectively. The list of all detected proteins is provided in Supplementary Table 5. **f** and **g)** Transduction efficiency of the PLE indicated upon infection by ΔCRISPR-Cas ICP1. (f) PLE11 mutants relative to the wild type (WT) control. (g) PLE1 and PLE4 Δ*tmp*-mutant derivatives with an empty vector (EV) or a plasmid expressing Rta normalized to the WT control with an EV. The bar represents the mean; each dot represents a biological replicate; the error bars indicate the standard deviation, and DL indicates the detection limit. Statistical analyses are the results of Student’s t-test by column pair (ns p > 0.05, *p < 0.05, **p < 0.01, ****p < 0.0001). **h)** Model for PLE-mediated tail manipulation and Rta-mediated anti-phage defense. All PLEs encode an alternative tape measure protein that is selectively incorporated into PLE virions through an unknown and likely conserved mechanism. PLE11 uniquely encodes Rta, which is not necessary for the assembly of chimeric tails but protects *V. cholerae* from infection by ICP1 possessing anti-PLE mechanisms like CRISPR-Cas that would typically circumvent all other known PLE-encoded anti-phage inhibitors via degradation of the PLE genome.

As ICP1’s TMP is the apparent target of Rta, we hypothesized that PLE11 may achieve tail formation in the presence of Rta by encoding an alternative, Rta-resistant TMP of its own. Although no other satellites have been reported to encode a TMP, we found that PLE11 encodes a protein with similarity to probable TMPs using HHpred (Fig. 4c). Further, bioinformatic and structural analyses of neighboring genes revealed that PLE11 additionally encodes a putative tail assembly chaperone (TAC) and baseplate hub protein (BhuB) (Fig. 4c and Extended Data Fig. 6). Since assembly of the tail tube around the TMP initiates from a multi-component baseplate complex and requires a stabilizing TAC^17^, these results indicate that PLE11 encodes a partial tail assembly pathway. Intriguingly, although Rta is unique to PLE11, *tmp* and *tac* genes are conserved across PLE variants, clustering into two sequence groups (exemplified by PLE11 and PLE4, Fig. 4c and Extended Data Fig. 6). Based on this, we hypothesized that PLE11 assembles chimeric tails by hijacking ICP1 components (e.g., tail tube, sheath, baseplate components, and tail fibers) while substituting the TMP, TAC, and BhuB with PLE-encoded proteins (which all have 20%, 20% and 41% identity to ICP1-encoded counterparts, respectively (Supplementary Table 10)), thus evading Rta activity.

To test this hypothesis, we investigated the structural and compositional differences between ICP1 and PLE11 virions. Since the TMP determines tail length^17^ and the PLE11 TMP and ICP1 TMP differ by 95 amino acids, we expected PLE TMP incorporation to result in shorter tails. Examination of purified ICP1 and PLE virions by TEM confirmed the expected 10nm difference between phage and PLE tails (Fig. 4b). Western blot analysis of purified virions confirmed the incorporation of PLE-encoded TMP into PLE11 virions, with exclusion of ICP1’s TMP and BhuB (Fig. 4d). To further validate these findings and provide a more thorough evaluation of protein composition, we analyzed the virions by mass spectrometry. For ICP1, we detected all major predicted proteins comprising the capsid and the tail (Fig. 4f, Supplementary Table 5). In PLE11 virions, we detected ICP1-encoded capsid proteins as expected, as well as most of ICP1’s tail components, with the notable exception of ICP1’s TMP and BhuB. Accordingly, these analyses demonstrated the incorporation of PLE11’s TMP and BhuB in PLE11 virions (Fig. 4e, Supplementary Table 5). Two additional PLE proteins without predicted functions were also found in the virions (Supplementary Table 5). Collectively, these data demonstrate that PLE11 uses both satellite- and phage-derived proteins to assemble functional chimeric tails, representing a novel mechanism of protein piracy and manipulation among phage satellites that has not been observed previously.

The substitution of PLE11’s TMP in the tails of virions provides a plausible mechanism for how PLE11 escapes the inhibition of phage tail assembly imposed by Rta’s targeting of the ICP1’s TMP. We inferred that PLE11’s TMP would thus be essential for its horizontal transduction when Rta was restricting ICP1’s TMP. To test this, we compared the transduction efficiency of several PLE11 mutant derivatives to that of the wild type element. In line with our hypothesis, the deletion of *tmp* in PLE11 caused a dramatic ∼99.9% reduction in transduction efficiency, confirming its essential role (Fig 4f). In a Δ*rta*Δ*tmp* double mutant, transduction was restored significantly, indicating that relief of Rta-mediated restriction of the phage’s TMP allows the incorporation of ICP1’s TMP into PLE virions. Complementing Δ*rta*Δ*tmp* with ectopic expression of Rta exacerbated the Δ*rta*Δ*tmp* transduction defect (Extended Data Fig. 7), further supporting this model. However, the transduction efficiency of the double Δ*rta*Δ*tmp* mutant remained markedly below wild type levels (Fig 4f), suggesting that PLE11 cannot efficiently substitute ICP1’s TMP even in the absence of Rta.

Intriguingly, the deletion of *rta* alone had no effect on PLE11 transduction (Fig. 4f), indicating Rta’s interference with ICP1’s tail assembly is not required for incorporating the PLE TMP. Consistent with this, all PLEs encode *tmp* and *tac* genes, suggesting they can produce hybrid tails wherein the PLE-encoded TMP is incorporated. Rta is not part of the PLE tail gene cluster and is uniquely found in PLE11; thus, it is not likely a factor required for the manipulation of tail assembly. In parallel with observations from PLE11, Δ*tmp* derivatives of PLE1 and PLE4, which represent distinct *tmp*-*tac* gene clusters (Fig. 4c), showed significant transduction defects (96% and 99% reduced efficiency compared to wild type, respectively) (Fig. 4g), suggesting manipulation of tail assembly to incorporate the PLE TMP is a broad feature of this family of satellites. Ectopic Rta expression had no effect on wild type PLE1 or PLE4 transduction, consistent with these PLEs using their TMP for virion assembly and thus being impervious to Rta’s restriction of ICP1’s TMP. However, like PLE11, Rta exacerbated the transduction defects of PLE1 and PLE4 Δ*tmp* mutants (Fig. 4g), consistent with the model that PLEs can less efficiently use ICP1’s TMP and, when forced to do so, are sensitized to Rta-mediated restriction.

Together, these findings establish that PLEs are the first family of phage satellites known to manipulate viral tail assembly to incorporate satellite- and phage-encoded structural components into chimeric tails. The unique adaptation of Rta encoded by the novel PLE variant, PLE11, capitalizes on an already existing, but yet unknown, pathway to manipulate phage tail assembly (Fig. 4h). Importantly, Rta’s activity provides robust inhibition of ICP1’s tail formation even when the PLE genome is targeted for degradation by ICP1’s nucleolytic anti-PLE mechanisms (Fig. 4h), thus providing a selective advantage to PLE11(+) *V. cholerae* in the face of CRISPR-Cas(+) or Adi(+) phages. These mechanistic insights help to explain the selective sweep of PLE11(+) BD-1.2, sublineage strains during this surveillance period, and the prolonged period with which PLE11(+) *V. cholerae* strains went unchallenged by ICP1.

## Discussion

The interplay between anti-phage defenses and the MGEs that encode them has proven far more nuanced than initially envisioned. Phage satellites, for instance, have been shown to shield bacterial populations from phages directly through co-opting a phage’s resources that ultimately permit satellite spread or by encoding broad-acting phage defense cargo that is auxiliary to the parasite’s horizontal transmission^20^. This former anti-phage activity is typically narrow spectrum, relying on precise molecular interactions between cognate phage and satellite factors. In two ways, our data contribute to an expanded perspective of how satellites defend bacterial populations by manipulating virion assembly and altruistic phage defense. First, it is well documented that phage satellites manipulate capsid assembly and genome packaging^21,22^, but PLEs’ manipulation of tail assembly to produce chimeric tails reveals a previously unknown aspect of satellite-mediated reprogramming of virion assembly. While further work is necessary to decipher the benefits of modulating tail assembly, intriguingly, putative satellites found in other *Vibrio* species and unrelated satellites found outside of the *Vibrionaceae* also encode predicted *tmp* genes (Extended Data Fig. 8), suggesting that manipulating tail assembly is a broader facet of satellite biology. Second, PLE11’s Rta provides the first example of a satellite-encoded defense protein that protects the bacterial population by interfering with the assembly of the parasitized phage independent of satellite transmission. Conversely, other satellites have been shown to encode broad-acting phage defenses that spare the phages they parasitize^20^, a strategy that presumably benefits both the satellite and the parasitized phage by limiting phage predation of the shared bacterial host. For Rta-mediated defense that restricts phage assembly under circumstances where the PLE genome is degraded, phage defense is altruistic as it eliminates transmission of both the parasitized phage and the satellite (Fig. 4h). This seemingly selfless defense could be favorable given the clonal populations typical of cholera epidemics, wherein limiting phage predation still ensures vertical transmission of the satellite within the population.

Our results indicate that all PLEs encode Rta-independent mechanism(s) to disrupt phage tail assembly. Although the mechanism(s) remains enigmatic, we speculate that the highly conserved regulatory small RNA, SviR, encoded by all PLEs contributes to this phenomenon. SviR in the context of PLE1 was shown to bind to transcripts for ICP1’s TAC (*gp82*) and BhuB (*gp78*)^23^, suggesting it may contribute to decreasing the abundance of these phage factors, favoring chimeric tail assembly and inhibiting phage tail assembly. In support of this prediction, SviR was shown to downregulate levels of ICP1’s capsid morphogenesis factor Gp120 during infection of PLE1(+) *V. cholerae*^23^, and notably, Gp120 is absent from PLE11 particles (Supplementary Table 5). Interestingly, not all PLEs encode an alternative baseplate hub, suggesting that even within this satellite family, PLEs have evolved multiple strategies to manipulate tail assembly. Satellites’ hijacking of phage proteins has provided many examples of distinct mechanisms to disrupt virion assembly of their cognate parasitized phage, and we expect this to continue to expand with the growing appreciation of satellite diversity and abundance^24^.

Notably, Rta is unique among PLE-encoded factors that interfere with ICP1’s lifecycle in its ability to restrict phage assembly even under circumstances where the PLE genome is degraded. Whether such potent inhibition by Rta in the face of nucleolytic degradation is attributed to high expression levels or early timing of gene expression, prolonged protein stability, or the nature of its targeting of ICP1’s TMP remains to be elucidated, and the lack of recognizable bioinformatic signatures of Rta hinders predictions. The molecular diversity of phage defense systems is striking from the more than one hundred defense systems discovered to date^1^. However, the targeting of tail assembly as a defense strategy has only very recently been discovered^25–27^, leaving our understanding of these defense mechanisms in its early stages. With Rta as the fourth example, it will be interesting for future works to explore the mechanistic comparisons between tail-targeting defenses. Furthermore, examining the convergent evolution of diverse satellites to incorporate such inhibitors into their defense arsenal while, in some cases, developing strategies for manipulation of tail assembly will also be informative.

In Bangladesh, the observed oscillations in the dominance of BD-1 and BD-2 lineages of *V. cholerae*^15^ and ICP1 genotypes with unique repertoires of anti-PLE counter-defenses^13^ are consistent with negative frequency-dependent selection dynamics and point to the maintenance of genetic diversity in under-sampled reservoirs. Such reservoirs may include asymptomatically infected individuals and/or the aquatic environment, though the precise location(s) and degree of diversity within such a reservoir remain unknown. While these oscillations provide a framework for further elucidation of the evolution of 7PET *V. cholerae* and its phages, it is important to recognize that Bangladesh is part of a wider transmission network for cholera across this region, and surveillance of phages in cholera stool samples is nonstandard, making widespread dynamics challenging to predict.

Anti-phage defense systems are seldom studied beyond simplified laboratory conditions, with few mechanisms investigated in their native hosts. This study provides conclusive evidence that predation by the lytic phage ICP1 drives the selection of 7PET *V. cholerae*. The acquisition of PLE11 into the BD-1.2 sublineage enabled these strains to resist contemporaneous ICP1 phages, facilitating their proliferation and likely contributing to the unusually devastating cholera outbreak in the spring of 2022^15^. Our results provide the missing mechanistic evidence supporting previous accounts of the assumed role of phage predation in limiting the duration and severity of cholera epidemics^28–30^. Additionally, our surveillance highlights the ongoing evolution of *V. cholerae* in the Bay of Bengal, emphasizing the importance of sustained genomic and phenotypic monitoring of both phages and bacteria to track the emergence and spread of 7PET *V. cholerae*. Beyond SNP-based phylogenetic analyses of the core genome, our findings underscore the necessity of examining variable accessory gene content to understand the success of emerging lineages and their potential to seed global cholera epidemics. Moving forward, deviations from expected patterns of phage susceptibility could inform public health measures, with phage-resistant linages prioritized as variants of concern. Importantly, the coevolution of ICP1 to counter bacterial defenses highlights the necessity of integrating mechanistic insight of circulating phage genotypes when developing these responses.

## Methods

### Statistics & Data reproducibility

Where applicable, statistical analyses were conducted using unpaired, two-tailed Student’s t-tests in GraphPad Prism. Relevant statistical results, including p-values and standard deviations, are reported in the figure legends alongside the data.

### Collection of patient stool samples and isolation of bacteria and phage

Cholera patient stool samples were collected and screened for *V. cholerae* and phages as described previously^31^. Stool samples were collected from suspected cholera patients at the icddr,b Dhaka Hospital and the Government Health Complex in Mathbaria, Pirojpur, under protocol number PR-16083 approved by the icddr,b Ethical Review Committee, with written consent obtained from participants or their guardians. Samples positive for *V. cholerae* serogroup O1 and/or O139 on VC RapidDipStick test (RDT; Span Diagnostics, Surat, India) were de-identified and stored at −80°C with w/v 20% glycerol. For on-site isolation of *V. cholerae*, positive samples were enriched in alkaline peptone water (APW, pH 8.4, Difco) at 37°C for 6–8 hours and then cultured overnight on taurocholate tellurite-gelatin agar (TTGA, Difco). *V. cholerae* appearing colonies were further confirmed using previously described biochemical and serological methods^32^. Additional purification of *V. cholerae* and isolation of phages from stool was performed at the University of California, Berkeley. *V. cholerae* isolates were further purified twice on LB agar plates and analyzed by PCR for PLE and/or whole-genome sequencing (see Supplementary Table 7 for primers). For phage isolation, a panel of *V. cholerae* hosts (including PLE(-) E7946 and PLE11(+) BFS783) was used to probe for phages from stool. Where possible, phages were isolated and purified on the cognate *V. cholerae* strain isolated from the stool sample. Bacterial hosts were grown to the mid-log phase, incubated with a small amount of frozen stool sample collected on a pipette tip (and dilutions thereof), and the mixture was plated in 0.5% LB top agar. Single plaques were picked and purified twice on the same host before being analyzed by PCR for *adi*, CRISPR-*cas*/*odn*, or TMP mutations using the primers listed in Supplementary Table 7 and/or whole genome sequencing.

### Whole genome sequencing

Genomic DNA from phages and bacteria was purified using Monarch Genomic DNA Purification Kit (New England BioLabs). Phage samples were initially treated with DNase I to remove non-encapsidated DNA. Illumina sequencing (150- by 150-bp paired end) was performed by the Microbial Genome Sequencing Center or SeqCenter (for all bacteria and most phage), and Nanopore sequencing was performed by the Barker Sequencing Core at the University of California, Berkeley (for a subset of phage isolates). Genomes were assembled using SPAdes^33^, and for escape phages selected on PLE11(+) *V. cholerae*, genomes were analyzed using BreSeq (v0.33)^34^.

### Bioinformatic analysis

The PLE genomes were aligned based on gene product identity using clinker^35^ at a 30% identity cutoff. We performed BLASTn searches against ICP1 genomes using *odn*, *adi*, and CRISPR-cas from ICP1_2001_Dha_0, ICP1_2006_Dha_E or ICP1_2011_Dha_A as queries. TMP substitutions were called if the sequence differed from ICP1_2006_Dha_E or ICP1_2011_Dha_A. ICP1 CRISPR spacers were manually annotated between direct repeats in the CRISPR arrays. The phage phylogeny was built by comparing whole genome sequences of 29 phages isolated from this study and 67 isolates from previous work^6^ using tBLASTx analysis from ViPTree^36^. ICP1 and PLE11 gene products identified in proteomics were analyzed for functional predictions using HHPred^37^ or extracted from previous work (for ICP1)^6^.

The phylogeny of bacterial genomes was calculated as described previously^15^. Briefly, fastp v0.23.2^38^ was used to evaluate the quality of the raw shotgun paired-end sequences. Genetic variants were identified by mapping the raw reads to the *V. cholerae* N16961 reference genome (NCBI accession IDs: NC_002505.1 and NC_002506.1) using snippy v4.6.0^39^. Phylogenetic analysis was performed using IQ-TREE v2.2.0^40^ with 1000 bootstrap and the best fitted evolutionary model was selected using ModelFinder^41^. Spades v3.15.4 genome assembler was used to generate contigs. Each of the 10 previously known PLEs^13^ and PLE11 were used as BLASTn queries against the *V. cholerae* genomes and annotated in the phylogeny. Lists of the SNPs in the core genome and strains used to build the phylogeny are in Supplementary Tables 8 and 9, respectively.

The structural predictions for TAC^PLE4^ and Rta were made using ColabFold^42^ on COSMIC2^43^ and GoogleColab Structural alignments TAC^HK97^ (PDB 2OB9), TAC^PLE1^ (5IR0), and predicted TAC^PLE4^ were done on ChimeraX^44^ using “Smith-Watermann ssFraction 0.8008 matrix BLOSUM-45 hgap 10 sgap 10 ogap 4” parameters. RMSD values for aligned pruned amino acid residues are reported. Putative satellite genomes from non-cholera *Vibrio* spp.,^31^ and cf-PICIs^45^ were analyzed for the presence of tape measure proteins and integrases using BLASTp and HHPred^37^. Genome visualizations were generated with R, utilizing the gggenes package.

### Bacterial growth and cloning

Bacterial and phage strains are reported in Supplementary Tables 1, 2, and 6. *V. cholerae* cultures were grown in LB broth at 37°C with aeration or on LB agar plates. When required, antibiotics were used at the following concentrations: 100 µg/mL spectinomycin (Spec), 75 µg/mL kanamycin (Kan), 2.5 µg/mL chloramphenicol (Cm) in LB plates, and 1.25 µg/mL Cm in liquid cultures. *Escherichia coli* cultures were grown in LB broth at 37°C with aeration or on LB agar plates, and 25 µg/mL Cm was when needed. *V. cholerae* strains were made naturally competent through previously reported methods^46^ and then were transformed with DNA fragments generated from purified PCR products. To create the laboratory strain of PLE11(+) *V. cholerae,* a *kan^R^* cassette was inserted downstream of the last PLE11 ORF via the natural transformation of clinical strain BFS783. The PLE11:*kan^R^* was then transduced into *V. cholerae* E7496. Gene deletions in the PLE11 *kan^R^*E7946 derivative were made by replacing the target gene(s) with a *frt-spec^R^-frt* cassette by natural transformation, and where indicated, the *spec^R^*was cured as described previously^10^. To generate the plasmid p-*rta*, the *rta* coding sequence from PLE11 was cloned into the pKL06.2 vector using Gibson assembly and selected in *E. coli* XL-1 Blue. The purified plasmid was electroporated into *E. coli* S17 and conjugated into *V. cholerae* strains. All deletions and plasmid constructs were confirmed by PCR and Sanger sequencing.

### Phage propagation, generation of escapes, and phage engineering

For standard plaque assays, phage isolates were propagated on *V. cholerae* to generate confluent lysis plates, collected in STE buffer (100 mM NaCl, 10 mM Tris-Cl pH 8.0, 1 mM EDTA), treated with chloroform, and clarified by centrifugation at 5000 × g for 15 minutes. The aqueous layer containing phages was stored at 4°C. Escape phages were generated by picking individual plaques from plaque assays with the parental phage, ICP1_2006_Dha_E (for CRISPR(+)) or ICP1_2011_Dha_A (deleted of CRISPR spacers targeting PLE11 (ACMphi232), for Adi(+)), on PLE11(*+*) *V. cholerae* (JEF42). The plaques were purified three times, and then genomic DNA was extracted from high-titer stocks and whole-genome sequenced (ICP1_2006_Dha_E), or the *tmp* was analyzed by Sanger sequencing (ICP1_2011_Dha_A derivatives). Phages with engineered mutations were generated by passaging the parental phage on a *V. cholerae* strain expressing an inducible CRISPR-Cas system with a targeting spacer and a repair template encoding the desired mutation as described previously^47^. Individual plaques were purified, and the *tmp* sequence was confirmed by Sanger sequencing.

### Phage assays

Saturated *V. cholerae* cultures (OD_600_ >2) were diluted to OD_600_ = 0.05 and grown to OD_600_ between 0.3 and 0.5 (mid-log) at 37°C with aeration. Cultures were induced with 1 mM IPTG and/or 1.5 mM theophylline at OD_600_ = 0.2 when necessary. For spot assays, 200 µL or 300 µL mid-log *V. cholerae* cultures were mixed with 0.5% LB top agar (with required antibiotics and inducers, as needed) and poured on 100 mm or 150 mm plates, respectively. Next, 4 mL or 12 mL of this mix was poured onto 100 mm or 150 mm LB agar plates, respectively. Ten-fold serial dilutions of phage were prepared, and 3 µL of each dilution was spotted onto the prepared top agar with bacteria, allowed to dry, and incubated at 37°C for 6 hours. To quantify plaquing efficiencies, 10 µL of phage dilutions were mixed with 50 µL mid-log *V. cholerae* for 7-10 minutes at room temperature. The phage-bacteria mix was added to 0.5% LB top agar (4 mL) in 60 mm plates. Inducers and antibiotics were added when required. The plates were incubated at 37°C for 6-7 hours and then at room temperature overnight. The efficiency of plaquing was calculated from three biological replicates relative to the permissive empty vector control (for p-*rta*) or permissive PLE(-) control (for PLE11(+)). Data presented as images represent one of the three biological replicates, and additional replicates are reported in the extended data or supplementary information.

### Quantitative Polymerase Chain Reaction (qPCR)

PLE11 replication PCRs were performed as described^12^. Briefly, saturated *V. cholerae* cultures were diluted to OD_600_ = 0.05 and grown to OD_600_ = 0.3 at 37¼C with aeration. Immediately before infection (t=0), 100 µL samples were collected and boiled. The remaining culture was infected with phage (ICP1_2006_Dha_EΔCRISPR*Δcas2-3* or wild type ICP1_2006_Dha_E) at an MOI of 2.5, incubated at 37¼C with aeration for 20 minutes (t=20), collected and boiled. PLE11 DNA from boiled samples was measured in technical duplicates by quantitative PCR relative to a standard curve using primers indicated in Supplementary Table 7. The fold change in genome copy was calculated as the amount of PLE11 genome at t=20 relative to the PLE11 genome at t=0. qPCR experiments were performed with three biological replicates.

### Propagation and purification of ICP1 progeny on *V. cholerae* with p-*rta*

ICP1 production in the presence of an inducible PLE gene was done as described previously^10^. An overnight culture of *V. cholerae* strain with either p-*rta* or pEV was back diluted to OD_600_ = 0.05 in 50 mL LB broth with 1.25 µg/mL Cm and incubated at 37¼C with aeration. At OD_600_ = 0.2, 1 mM IPTG and 1.5 mM theophylline were added for induction, or an equivalent volume of sterile water was added in the uninduced control. Cultures were grown to OD_600_=0.3. and then infected with ICP1_2006_Dha_E_ΔCRISPR*Δcas2-3* at an MOI of 2.5. Upon lysis (∼90 mins), the culture was treated with 0.25 units/ml Benzonase and 10% chloroform for 5 minutes with shaking, centrifuged at 5,000 x g for 15 minutes at 4°C. The supernatant was centrifuged at 26,000 x g for 90 minutes at 4°C. The resulting phage pellet was resuspended in phage buffer (50 mM Tris–HCl pH 8.0, 100 mM NaCl, 10 mM MgSO_4_, 1 mM CaCl_2_) by rocking overnight at 4°C, treated with chloroform (1:1) and the aqueous layer was collected for further analysis. For TEM analysis of ICP1 propagated in the presence of p-*rta* +/-inducer (Fig. 2d), one biological replicate was performed; three biological replicates of TEM analysis of ICP1 propagated in the presence of an induced empty vector or induced p-*rta* are presented in Extended Data Fig. 3.

### Purification of ICP1 and PLE11 virions

ICP1 and PLE11 virions were generated by infecting 200 mL mid-log culture of PLE(-) *V. cholerae* and PLE11(+) *V. cholerae,* respectively, at MOI of 2.5 with ICP1_2006_Dha_EΔCRISPR*Δcas2-3*. After culture lysis, 0.25 units/mL Benzonase and 10 mL micellar mix (4 mL chloroform, 2 mL methanol, 25 mM sodium citrate, and 10 mM sodium deoxycholate) were added and mixed for 5 minutes at room temperature. The lysate was centrifuged at 5000 x g for 10 mins at 4°C; the aqueous supernatant was centrifuged at 26,000 x g for 90 minutes at 4°C. The supernatant was discarded, and the pellet was recovered in phage buffer by rocking overnight at 4°C. The resuspended pellet was chloroform treated (1:1), and the aqueous layer containing virions was collected. Next, to separate particles from aggregates and free proteins, the concentrated phage stock was subjected to bench-top gel filtration chromatography as follows: 3 g Bio-Rad P-10 Gel Fine 45-90 was hydrated in 50 mM Tris-Cl pH 8.0 at room temperature and then stored at 4°C until the next steps. BioRad Polyprep chromatography columns were packed with ∼3 mL of the hydrated gel to a 1.5-inch bed height. The gel was equilibrated with 15 mL phage buffer supplemented with Benzonase (0.25 units/mL) using gentle syringe pressure. The concentrated phage stock (∼400 µL) was loaded on the resin under gravity, followed by ∼3 mL of phage buffer with benzonase. The eluent was collected as 100 µL x 30 fractions, which were screened for virions using spot plate plaque assays for ICP1 and transductions for PLE11. The fractions with high titers were also visually inspected using TEM.

### Transmission electron microscopy

To stage the virions for imaging, copper mesh grids (Formvar/Carbon 300, Electron Microscopy Sciences) were loaded with 5 µL of samples for 1 minute, washed with 5 µL of sterile ddH_2_O for 15 seconds, and stained with 1% uranyl acetate for 30 seconds. The absorbent paper was used to wick liquids between each step gently. The grids were imaged on a Tecnai-12 electron microscope at 120 kV. Uncropped TEM images can be found in Supplementary Fig. 3.

### Particle measurements

The dimensions of the tails of ICP1 and PLE11 virions were measured using TEM images analyzed using Fiji^48^: the pixel distance was set to a scale distance (nm) using ‘Set Scale.’ Tail sizes were measured in a straight line from neck to base using the ‘Analyze’ > ‘Measure’ option. Particle measurements were performed on three biological replicates of particle purifications, for a total of n=170 particles measured for each ICP1 and PLE11.

### Mass spectrometry

Three of the highest concentration fractions from gel filtration were pooled (totaling 4.95E+12 TFU/mL for PLE11 virions and 3.00E+11 PFU/ml for ICP1 virions (ICP1_2006_Dha_EΔCRISPR*Δcas2-3*), and each pool was denatured in Lamelli buffer. 40 µL of each sample was run on Any-Kd Mini-PROTEAN TGX Precast gel (Bio-Rad). The gel was stained with GelCode Blue stain (ThermoFisher) for visualization, and the lanes with samples were cut into 1 mm^2^ pieces and destained. In-gel digestion and mass spectrometry analysis were conducted at the Vincent J. Coates Proteomics/Mass Spectrometry Laboratory (PMSL) at the University of California, Berkeley. Briefly, gel pieces were washed twice with 50% acetonitrile (ACN) and 50 mM ammonium bicarbonate for 15 minutes with shaking and then dehydrated with 100% ACN for 5 minutes, followed by air drying for 5 minutes. Pieces were further dried with 10 mM Tris(2-carboxyethyl)phosphine hydrochloride (TCEP) and 40 mM chloroacetamide (CAA) for 5 minutes at 70°C. The gel was rehydrated in 50 mM HEPES pH 8.0 in a minimal volume enough to cover the surface. For digestion, rehydrated pieces were incubated with 1 µg Trypsin (1:50 dilution) for 1 hour at room temperature, then supplemented with a minimal volume of 50 mM HEPES pH 8.0 and incubated overnight at 37°C. Peptides were extracted stepwise in treatment with 25% and 100% ACN. Peptide extracts were concentrated to 30-60 µL and acidified with formic acid.

The digested peptides were analyzed by online capillary nanoLC-MS/MS using a 25 cm reversed-phase column fabricated in-house (50 µm inner diameter, packed with ReproSil-Gold C18-1.9 μm resin, Dr. Maisch GmbH) equipped with laser-pulled nanoelectron spray emitter tip. Peptides were eluted at 100 nL/min on a Thermo Fisher Easy-nLC1200 using a 140 min linear gradient of 2– 40% buffer B (buffer A: 0.05% formic acid in water; buffer B: 0.05% formic acid in 95% acetonitrile in water). Peptides were ionized using a FLEX ion source and analyzed on a Fusion Lumos Tribrid Orbitrap Mass Spectrometer (Thermo Fisher Scientific), and data was acquired in Orbitrap mode with parameters as follows: MS1 resolution of 120,000 at 200 m/z and scan range of 350–1600 m/z. The top 20 most abundant ions were fragmented via collision-induced dissociation (CID) with 35% normalized collision energy, activation q of 0.25, and a two m/z precursor isolation width. Dynamic exclusion was set to a 30-second repeat duration, 20-second exclusion, and a single repeat count. RAW files were analyzed with PEAKS (Bioinformatics Solutions Inc.) using semi-specific cleavage at R (Arg) and K (Lys) (up to 4 missed cleavages), a precursor mass tolerance of 15 ppm, and fragment mass tolerance of 0.5 Da. Variable modifications included methionine oxidation, and cysteine carbamidomethylation was fixed. Peptide hits were filtered using a 1% FDR, with proteins requiring at least two unique peptides and 1% FDR. Label-free quantitation (LFQ) was performed with PEAKS using default parameters, except for selecting the top 2 peptides per protein with a minimum abundance of 10e4 and normalization based on TIC across technical replicates. Data presented in Fig. 4e are based on one biological replicate (note that the western blot analysis of particles was performed on a separate biological preparation of particles).

### Custom antibodies and western blotting

The antibody against the ICP1’s capsid protein (α-capsid^ICP1^) was used as described^49^. Polyclonal antibodies against ICP1’s Gp78 (α-BhuB^ICP1^), ICP1’s TMP (α-TMP^ICP1^), and PLE11’s TMP (α-TMP^PLE11^) were generated in rabbits by GenScript. Given the intrinsic toxicity and challenges in expressing tape measure proteins, regions of least relative disorder were predicted using the modeled structure generated by ColabFold^42^ and sequence-based predictions from DEPICTER2^50^. The codon-optimized DNA sequence for Met^252^-Ala^338^ from ICP1’s TMP and Met^205^-Asn^319^ from PLE11’s TMP were cloned in pET30a(+), 6x-His-tagged proteins were purified using Ni-affinity chromatography and used as antigens.

To analyze the presence of listed proteins in purified virions, samples were prepared in Lamelli buffer, denatured, and loaded onto SDS-PAGE in duplicate. The proteins were transferred to 0.22 PVDF membrane using TransBlot Turbo (BioRad) at 1.5V for 7 minutes, and the blot was blocked with 0.5% skim milk in TBST (20 mM TrisCl pH 7.6, 150 mM NaCl 0.1% Tween 20). The blot was cut at 50 kDa and 25 kDa. The >50kDa pieces were probed with α-TMP^ICP1^, and α-TMP^PLE11^ (diluted 1:1500), blots from 50 to 25 kDa were probed with α-capsid^ICP1^ (diluted 1:1500), and pieces <25 kDa were probed with α-BhuB^ICP1^(diluted 1:3000). The primary antibodies were diluted in TBST with 2% bovine serum albumin (BSA). Blots were incubated in primary antibodies overnight at 4°C, washed thrice in TBS (20 mM TrisCl pH 7.6, 150 mM NaCl), and incubated in goat anti-rabbit secondary antibodies in TBST with 2% BSA for 45 minutes, and washed twice in TBS. Blots were developed in ECL chemiluminescence reagent (BioRad) and imaged on BioRad ChemiDoc. Data presented in Fig. 4d are based on one biological replicate (note that the Mass spec analysis of particles was performed on a separate biological preparation of particles). Uncropped western blot images can be found in Supplementary Fig. 3.

### Transduction assays

Donor strains of *V. cholerae* with PLE marked with a *kan^R^* gene downstream of the last PLE gene were grown to saturation. The donor strains were back diluted to OD_600_= 0.05 in 2mL LB broth, grown to OD_600_ = 0.3 and infected with ICP1_2006_Dha_E_ΔCRISPR*Δcas2-3* at MOI of 2.5. After 5 minutes, the unbound phage was washed off, and the infected cells were resuspended in fresh media. After ∼20 mins, the lysates were collected, treated with 20 µL chloroform, and centrifuged at 5,000 x g for 15 minutes at 4°C. Where necessary, Cm 1.25 µg/mL was added to the media for the growth of donor strains but was excluded from the washing and resuspension steps. In parallel, the recipient strain, *V. cholerae (ΔlacZ:: spec*), was grown for 6-7 hours and supplemented with 10 mM MgSO_4_ right before adding the lysate for transductions. Next, 20µL of supernatant was mixed with 180 µL of the recipient. Ten-fold dilutions of the mix were prepared in LB broth and incubated at 37°C with shaking for 20 minutes. All dilutions were then plated on LB agar plates containing kanamycin and spectinomycin, and the resulting colonies were counted to calculate transducing-forming units per mL of the donor lysate. The transduction efficiency was calculated relative to wildtype PLE, wild type with empty vector, or empty vector controls as applicable. Transduction efficiency was calculated from three biological replicates.

## Data availability

Sequence data for *V. cholerae* and ICP1 isolates from clinical samples and for ICP1 escape phages have been deposited in the NCBI Sequence Reads Archive (SRA) under BioProject PRJNA1195958. The PLE11 sequence has been deposited to GenBank (accession PQ783903). The mass spectrometry proteomics data have been deposited to the ProteomeXchange Consortium via the PRIDE partner repository with the dataset identifier PXD058665 under the DOI 10.6019/PXD058665. Reviewer links were provided for submission, and data will be released upon publication.

## Code availability

No custom code was used.

## Acknowledgments

The authors are especially thankful for the support of the icddr,b hospital and the Molecular Ecology and Metagenomics Laboratory staff. The authors thank Dr. Angus Angermeyer for the initial bioinformatic investigation of PLE11 and members of the Seed lab for critical feedback and thoughtful discussion regarding this project. The authors thank the staff at the electron microscopy facility and the Vincent J. Coates Proteomics/Mass Spectrometry Laboratory Core Facility (RRID: SCR_025852) at the University of California Berkeley. The authors thank members of the Center for Structural Genomics of Infectious Diseases (CSGID) for determining the X-ray diffraction structure of the TAC from PLE1 (PDB:51R0). CSGID was funded with U.S. Federal funds from the National Institute of Allergy and Infectious Diseases, National Institutes of Health Department of Health and Human Services under Contract number HHSN272201700060C. This work was supported by the National Institute of Allergy and Infectious Diseases (grants R01AI127652 and R01AI153303 to K.D.S.). Its contents are solely the responsibility of the authors and do not necessarily represent the official views of the National Institute of Allergy and Infectious Diseases or NIH. icddr,b gratefully acknowledges the Government of the People’s Republic of Bangladesh and the Global Affairs Canada (GAC) for their unrestricted support.

## Author contributions

Y.M. & C.M.B.: Conceptualization, Investigation, Visualization, Formal analysis, Writing – Original draft preparation, Writing – Review & Editing. J.E.F.: Investigation. M.M.M.: Formal analysis. M.T.I. Resources. M.S.: Resources, Project Administration. T. A: Resources, Supervision. M.A.: Funding acquisition, Resources, Supervision. K.D.S.: Conceptualization, Investigation, Writing – Original draft preparation, Writing – Review & Editing, Project administration, Funding acquisition. All authors discussed the results, commented on and approved the final manuscript.

## Competing interests

The authors declare no competing interests

**Correspondence and requests for materials** should be addressed to Kimberley Seed.

