## Supplementary Figures for "Capturing dynamic phage-pathogen coevolution by clinical surveillance"

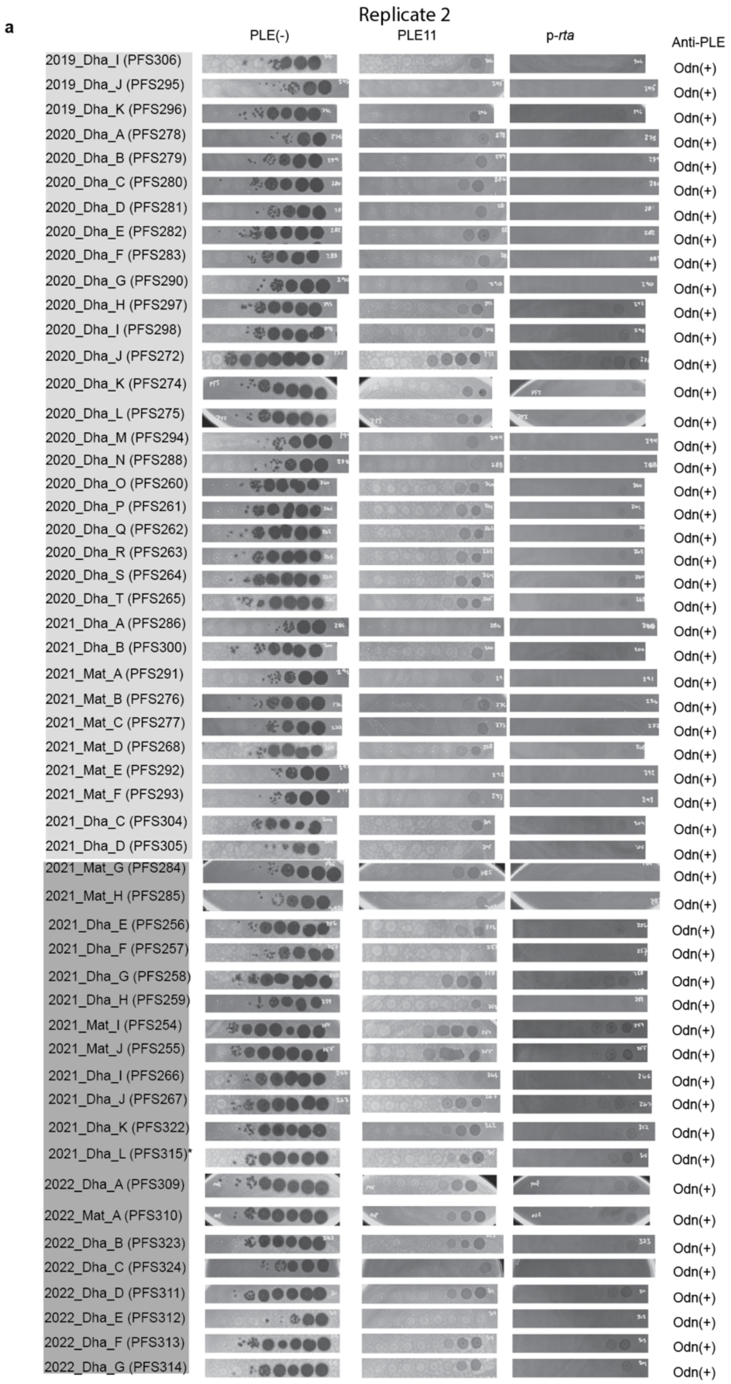

a
Replicate 2
PLE(-)
PLE11
p-rta
Anti-PLE
2019_Dha_I (PFS306)
2019_Dha_J (PFS295)
2019_Dha_K (PFS296)
2020_Dha_A (PFS278)
2020_Dha_B (PFS279)
2020_Dha_C (PFS280)
2020_Dha_D (PFS281)
2020_Dha_E (PFS282)
2020_Dha_F (PFS283)
2020_Dha_G (PFS290)
2020_Dha_H (PFS297)
2020_Dha_I (PFS298)
2020_Dha_J (PFS272)
2020_Dha_K (PFS274)
2020_Dha_L (PFS275)
2020_Dha_M (PFS294)
2020_Dha_N (PFS288)
2020_Dha_O (PFS260)
2020_Dha_P (PFS261)
2020_Dha_Q (PFS262)
2020_Dha_R (PFS263)
2020_Dha_S (PFS264)
2020_Dha_T (PFS265)
2021_Dha_A (PFS286)
2021_Dha_B (PFS300)
2021_Mat_A (PFS291)
2021_Mat_B (PFS276)
2021_Mat_C (PFS277)
2021_Mat_D (PFS268)
2021_Mat_E (PFS292)
2021_Mat_F (PFS293)
2021_Dha_C (PFS304)
2021_Dha_D (PFS305)
2021_Mat_G (PFS284)
2021_Mat_H (PFS285)
2021_Dha_E (PFS256)
2021_Dha_F (PFS257)
2021_Dha_G (PFS258)
2021_Dha_H (PFS259)
2021_Mat_I (PFS254)
2021_Mat_J (PFS255)
2021_Dha_I (PFS266)
2021_Dha_J (PFS267)
2021_Dha_K (PFS322)
2021_Dha_L (PFS315)*
2022_Dha_A (PFS309)
2022_Mat_A (PFS310)
2022_Dha_B (PFS323)
2022_Dha_C (PFS324)
2022_Dha_D (PFS311)
2022_Dha_E (PFS312)
2022_Dha_F (PFS313)
2022_Dha_G (PFS314)
Odn(+)

**a (continued)**

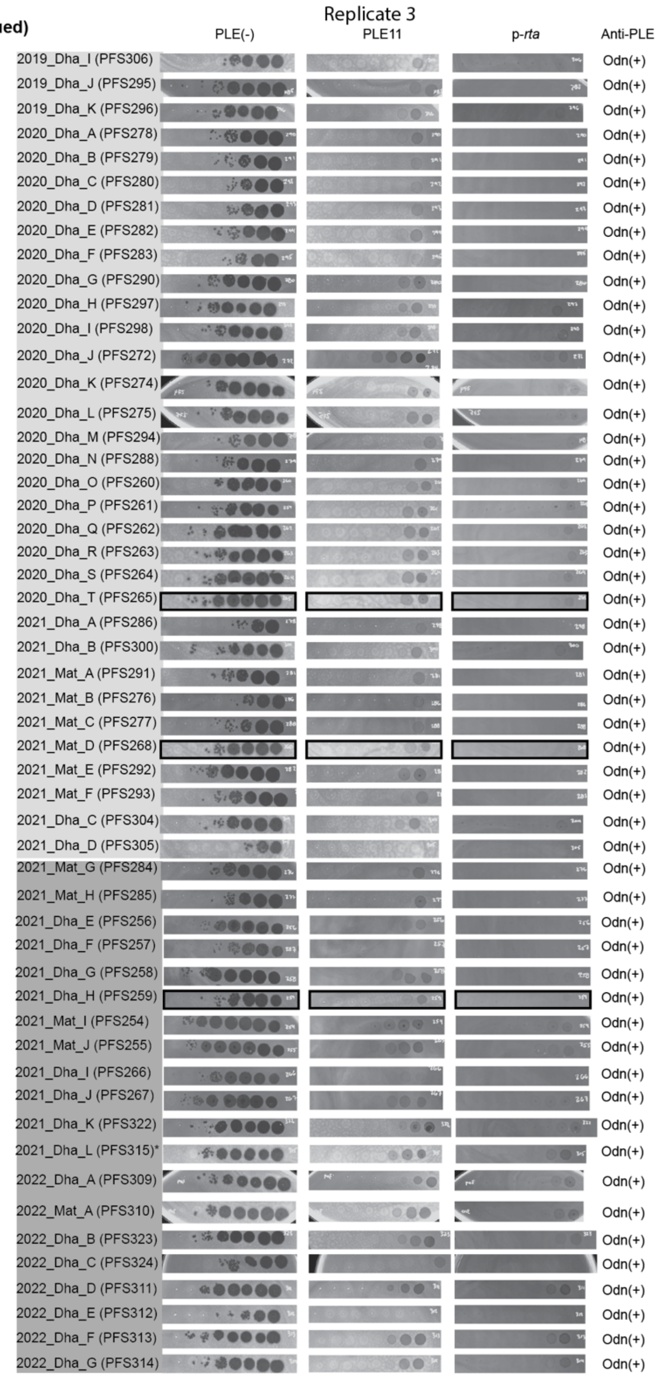

Replicate 3
PLE(-)
PLE11
p-rta
Anti-PLE
2019_Dha_I (PFS306) Odn(+)
2019_Dha_J (PFS295) Odn(+)
2019_Dha_K (PFS296) Odn(+)
2020_Dha_A (PFS278) Odn(+)
2020_Dha_B (PFS279) Odn(+)
2020_Dha_C (PFS280) Odn(+)
2020_Dha_D (PFS281) Odn(+)
2020_Dha_E (PFS282) Odn(+)
2020_Dha_F (PFS283) Odn(+)
2020_Dha_G (PFS290) Odn(+)
2020_Dha_H (PFS297) Odn(+)
2020_Dha_I (PFS298) Odn(+)
2020_Dha_J (PFS272) Odn(+)
2020_Dha_K (PFS274) Odn(+)
2020_Dha_L (PFS275) Odn(+)
2020_Dha_M (PFS294) Odn(+)
2020_Dha_N (PFS288) Odn(+)
2020_Dha_O (PFS260) Odn(+)
2020_Dha_P (PFS261) Odn(+)
2020_Dha_Q (PFS262) Odn(+)
2020_Dha_R (PFS263) Odn(+)
2020_Dha_S (PFS264) Odn(+)
2020_Dha_T (PFS265) Odn(+)
2021_Dha_A (PFS286) Odn(+)
2021_Dha_B (PFS300) Odn(+)
2021_Mat_A (PFS291) Odn(+)
2021_Mat_B (PFS276) Odn(+)
2021_Mat_C (PFS277) Odn(+)
2021_Mat_D (PFS268) Odn(+)
2021_Mat_E (PFS292) Odn(+)
2021_Mat_F (PFS293) Odn(+)
2021_Dha_C (PFS304) Odn(+)
2021_Dha_D (PFS305) Odn(+)
2021_Mat_G (PFS284) Odn(+)
2021_Mat_H (PFS285) Odn(+)
2021_Dha_E (PFS256) Odn(+)
2021_Dha_F (PFS257) Odn(+)
2021_Dha_G (PFS258) Odn(+)
2021_Dha_H (PFS259) Odn(+)
2021_Mat_I (PFS254) Odn(+)
2021_Mat_J (PFS255) Odn(+)
2021_Dha_I (PFS266) Odn(+)
2021_Dha_J (PFS267) Odn(+)
2021_Dha_K (PFS322) Odn(+)
2021_Dha_L (PFS315)* Odn(+)
2022_Dha_A (PFS309) Odn(+)
2022_Mat_A (PFS310) Odn(+)
2022_Dha_B (PFS323) Odn(+)
2022_Dha_C (PFS324) Odn(+)
2022_Dha_D (PFS311) Odn(+)
2022_Dha_E (PFS312) Odn(+)
2022_Dha_F (PFS313) Odn(+)
2022_Dha_G (PFS314) Odn(+)

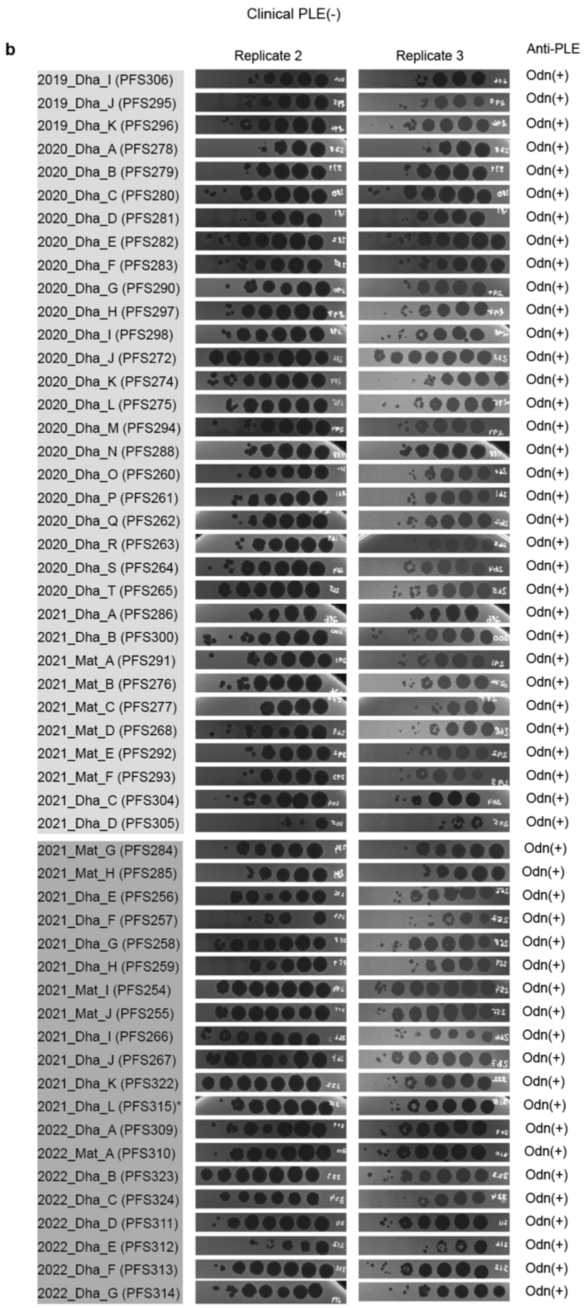


**Supplementary Fig. 1. PLE11 protects *V. cholerae* from pre- and co-circulating Odn(+) phage isolates.**

Biological replicates related to Extended Data Fig. 2. Plaquing of tenfold serially diluted ICP1 phage isolates from the pre-PLE11 and early-PLE11 periods (December 2019 to June 2022) on lawns of *V. cholerae* strain E7946, E7946 PLE11(+) and E7946 with a low-copy plasmid expressing PLE11 *rta* **(a)**, or on lawns of the clinical PLE(-) isolate BFS948 **(b)**. The gray background is the bacterial lawn, and the dark spots are zones of killing. Images that were used in the main text are indicated with black boxes. Standardized ICP1 names are shown (which include the year and location of isolation; Dha = Dhaka, Mat = Mathbaria), as well as lab designations (PFS#). The light gray shading indicates the period before PLE11 (pre-PLE11), the medium gray shading indicates the first year after the initial detection of PLE11 (early PLE11 period). All phage isolates from this period encode Odn as their sole anti-PLE mechanism; none of the whole genome sequenced isolates (n= 17) harbor substitutions in the TMP within the Rta-associated region (amino acid positions 314-387), but one isolate (marked by *) contains a TMP substitution (R132C) which does not provide resistance to PLE11 or Rta. Where isolates were genotyped by PCR for Odn vs CRISPR-Cas, the sequence of the *tmp* was not analyzed.

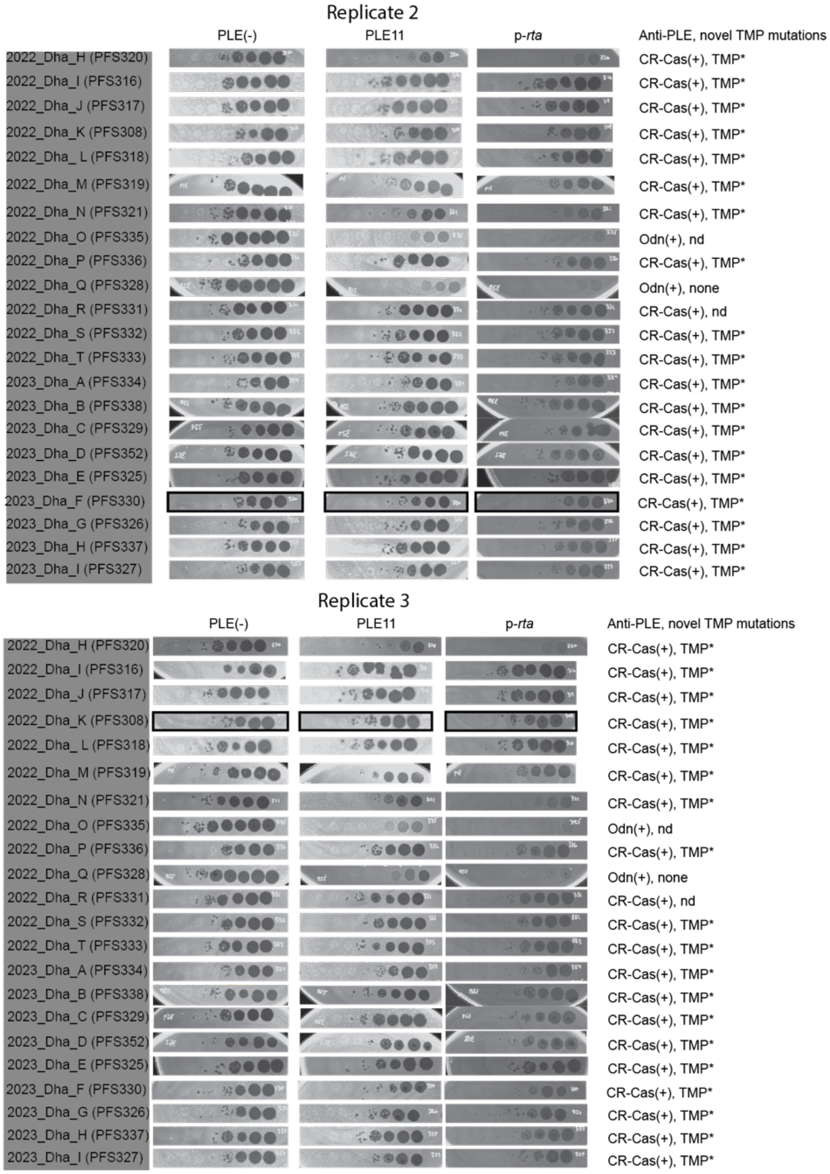


**Supplementary Figure 2. ICP1 isolates from the late-PLE11 period have evolved to overcome PLE11-mediated defense and Rta**

Biological replicates related to Extended Data Fig. 5. Plaquing of tenfold serially diluted ICP1 phage isolates from the late PLE11 period (July 2022 to December 2023) on lawns of *V. cholerae* strain E7946, E7946 PLE11(+) and E7946 with a low-copy plasmid expressing PLE11 *rta*. The gray background is the bacterial lawn, and the dark spots are zones of killing. Images that were used in the main text are indicated with black boxes. Standardized ICP1 names are shown (which include the year and location of isolation; Dha = Dhaka, Mat = Mathbaria), as well as lab designations (PFS#). The dark gray shading indicates the period after PLE11 (late-PLE11 period). Anti-PLE mechanisms and the presence of TMP mutations are indicated; TMP* refers to either substitution combination of L362P ± N355S, isolates for which the TMP was not analyzed are noted with “nd” for not-determined.

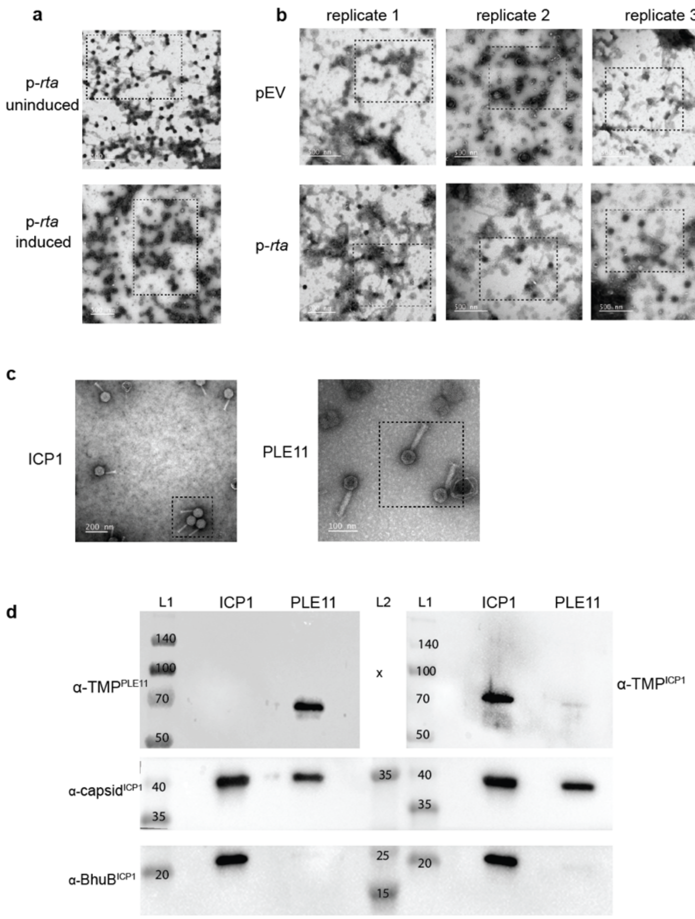


**Supplementary Fig. 3. Uncropped western blots and transmission electron micrographs.** **a-c)** Transmission electron micrographs (TEMs) used to generate data in Fig. 2d., Extended data Fig. 3, and Fig. 4a., respectively. Dashed boxes indicate the section of the image used in the main text. **d)** Uncropped western blots used to generate Fig. 4d. L1 and L2 indicate ladder 1 and ladder 2, respectively, and the size in kDa is noted on the blots. ICP1 and PLE indicate samples of purified virions used in this experiment. Antibodies used are indicated.
