## Extended data for "Capturing dynamic phage-pathogen coevolution by clinical surveillance"

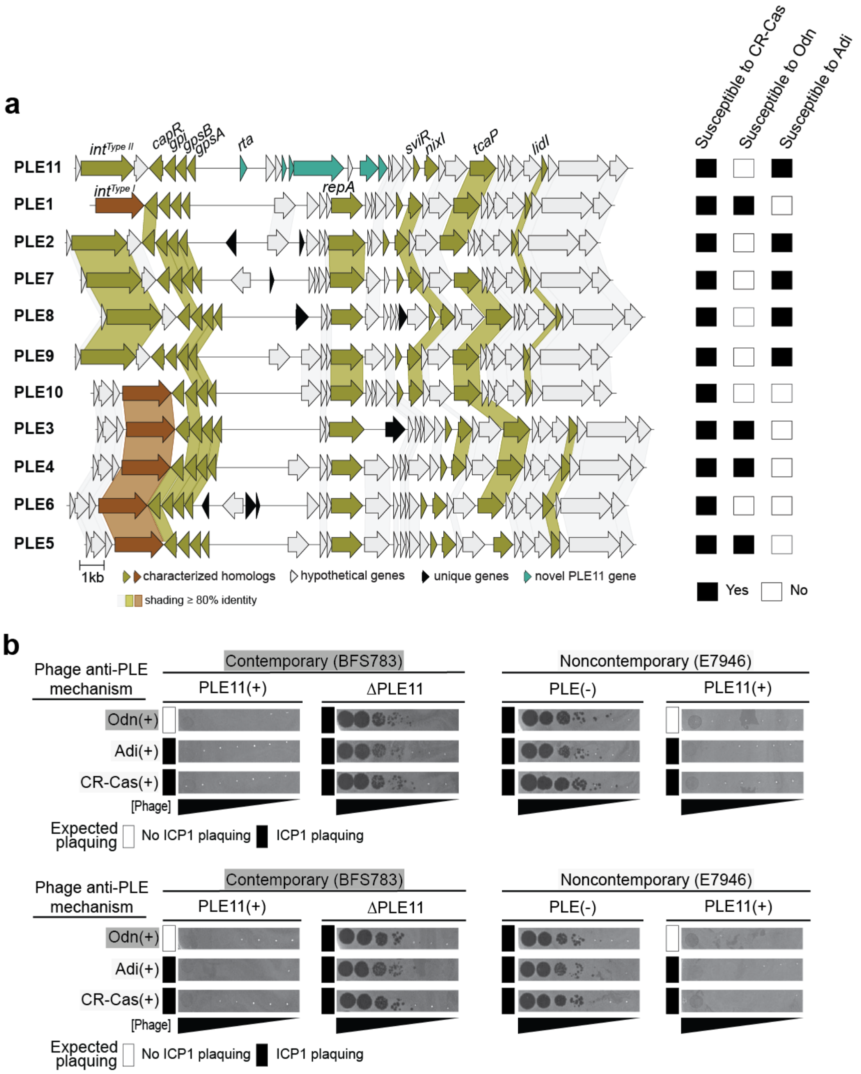


**Extended Data Fig. 1. The novel PLE variant, PLE11, inhibits ICP1 with known anti-PLE mechanisms**

**a)** Gene map comparison of *V. cholerae* PLEs. Genes were identified using tBLASTX and visualized using clinker. Similar genes encoding proteins in multiple PLEs have links drawn between them and are shaded if amino acid sequence identity >80%. Characterized genes from PLE1 and homologs in PLE11 are indicated; the Type II integrase was characterized in PLE2. The highly conserved regulatory RNA SviR (non-protein coding) is also shown. For each PLE, the expected susceptibility pattern to the known anti-PLE mechanisms is indicated (based on validated infection outcomes from characterized PLEs).

**b)** Replicates of spot plates for Fig. 1e. Plaquing of tenfold serially diluted ICP1 phage isolates with the anti-PLE mechanism indicated on lawns of *V. cholerae*. BFS783 refers to the first PLE11(+) clinical isolate identified in our surveillance (see arrow in Fig. 1c). E7946 PLE(-) and E7946 PLE11(+) are laboratory strains. The gray background is the bacterial lawn, and the dark spots are zones of killing. The expected plaquing phenotype (based on validated infection outcomes from characterized PLEs) is indicated for each bacterial host-phage pair. The plaquing phenotype for Odn(+) phage on the noncontemporary (E7946) strain is representative of all ICP1 isolates recovered during the pre-PLE11 and early PLE11 periods (n=53) (Extended Data Fig 2, Supplementary Fig. 1). CRISPR-Cas(+) and Adi(+) ICP1 isolates are historically collected isolates.

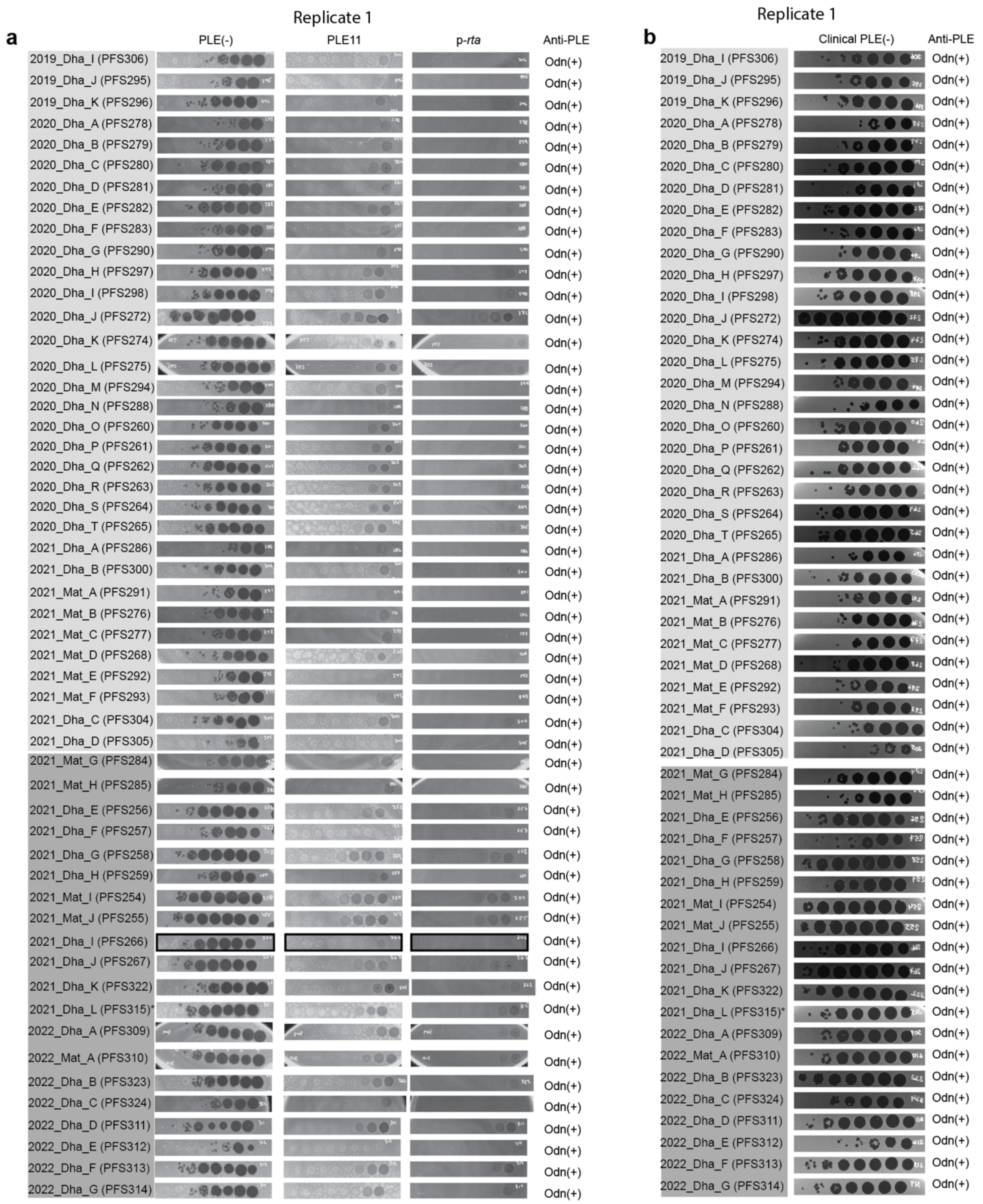


**Extended data Fig. 2. PLE11 protects *V. cholerae* from pre- and co-circulating Odn(+) phage isolates**

**a)** Plaquing of tenfold serially diluted ICP1 phage isolates from the pre-PLE11 and early-PLE11 periods (December 2019 to June 2022) on lawns of *V. cholerae* strain E7946, E7946 PLE11(+) and E7946 with a low-copy plasmid expressing PLE11 *rta*. Images used in the main text are indicated with black boxes.

**b)** Plaquing of tenfold serial diluted ICP1 phage isolates as in (a) on lawns of the clinical PLE(-) isolate BFS948. BFS948 was selected as the closest PLE(-) relative to the PLE11(+) isolate BFS783 used in Fig. 1. For both panels, standardized ICP1 names are shown (including the year and location of isolation: Dha = Dhaka, Mat = Mathbaria, and lab designations (PFS#)). The light gray shading indicates the period before PLE11 (pre-PLE11), and the medium gray shading indicates the first year after the initial detection of PLE11 (early PLE11 period). All phage isolates from this period encode Odn as their sole anti-PLE mechanism; none of the whole genome sequenced isolates (n= 17) harbor substitutions in the TMP within the Rta-associated region (amino acid positions 314-387), but one isolate (marked by *) contains a TMP substitution (R132C) which does not provide resistance to PLE11 or Rta. For isolates genotyped by PCR for Odn vs CRISPR-Cas, the sequence of the *tmp* was not analyzed. Additional biological replicates are included in Supplementary Fig. 1.

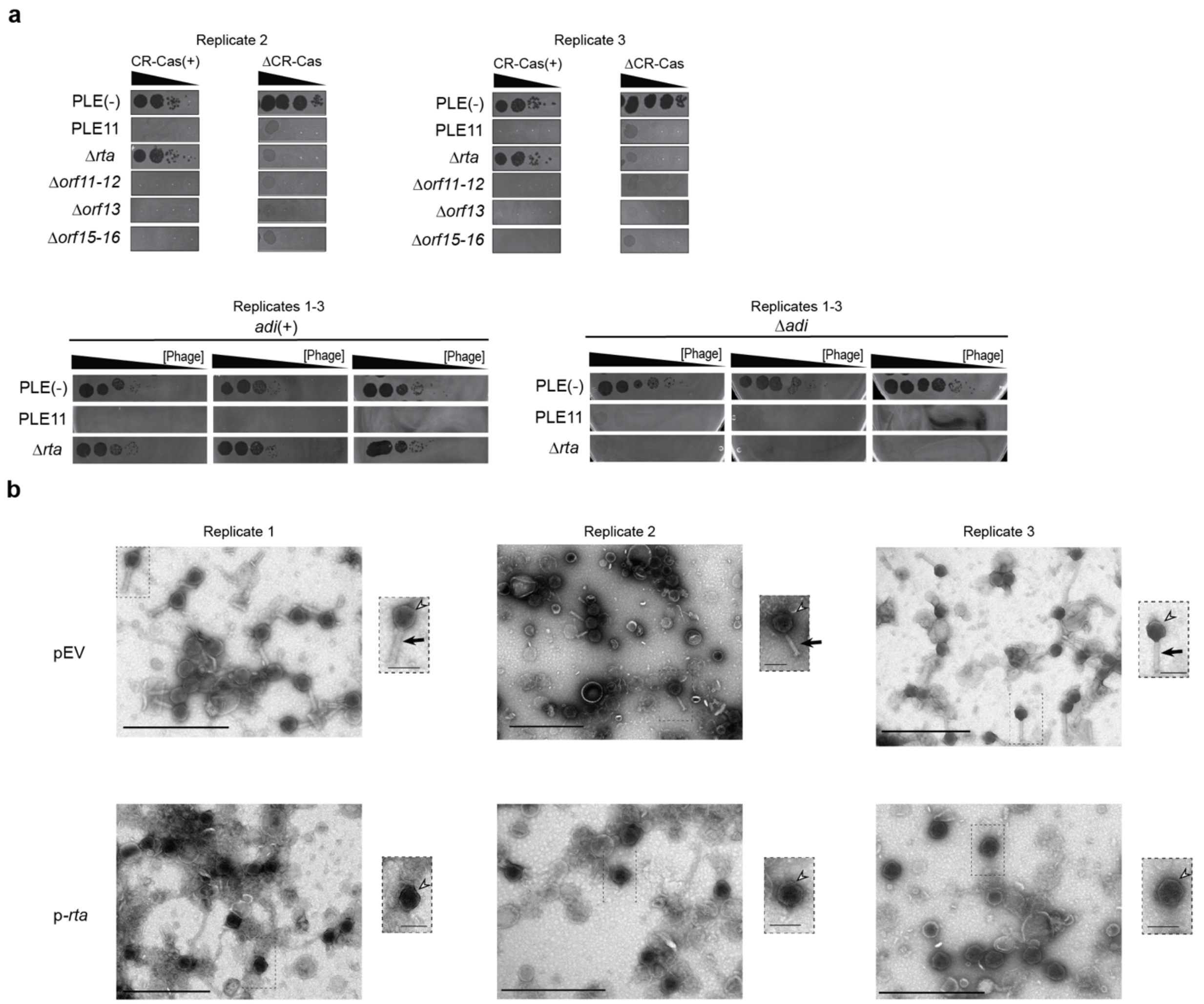


**Extended data Fig. 3. PLE11-encoded Rta restricts the assembly of tailed ICP1 virions**

**a)** Plaquing of tenfold serially diluted ICP1 CR-Cas(+/-) (top, replicates of Fig. 2a) or ICP1 Adi(+/-) (bottom) on lawns of *V. cholerae* strain E7946 and its PLE11(+) and PLE11 mutant derivatives. The gray background is the bacterial lawn, and the dark spots are zones of killing.

**b)** Representative transmission electron micrographs (TEMs) of particles produced following ICP1 infection of PLE(-) *V. cholerae* with Rta expressed from a low copy plasmid (p-*rta*) or an empty vector control (pEV)with inducer. The arrowheads indicate DNA-filled capsids, and the arrow indicates a tail. The scale bars are 500 nm and 100 nm for the zoomed-out and insets, respectively.

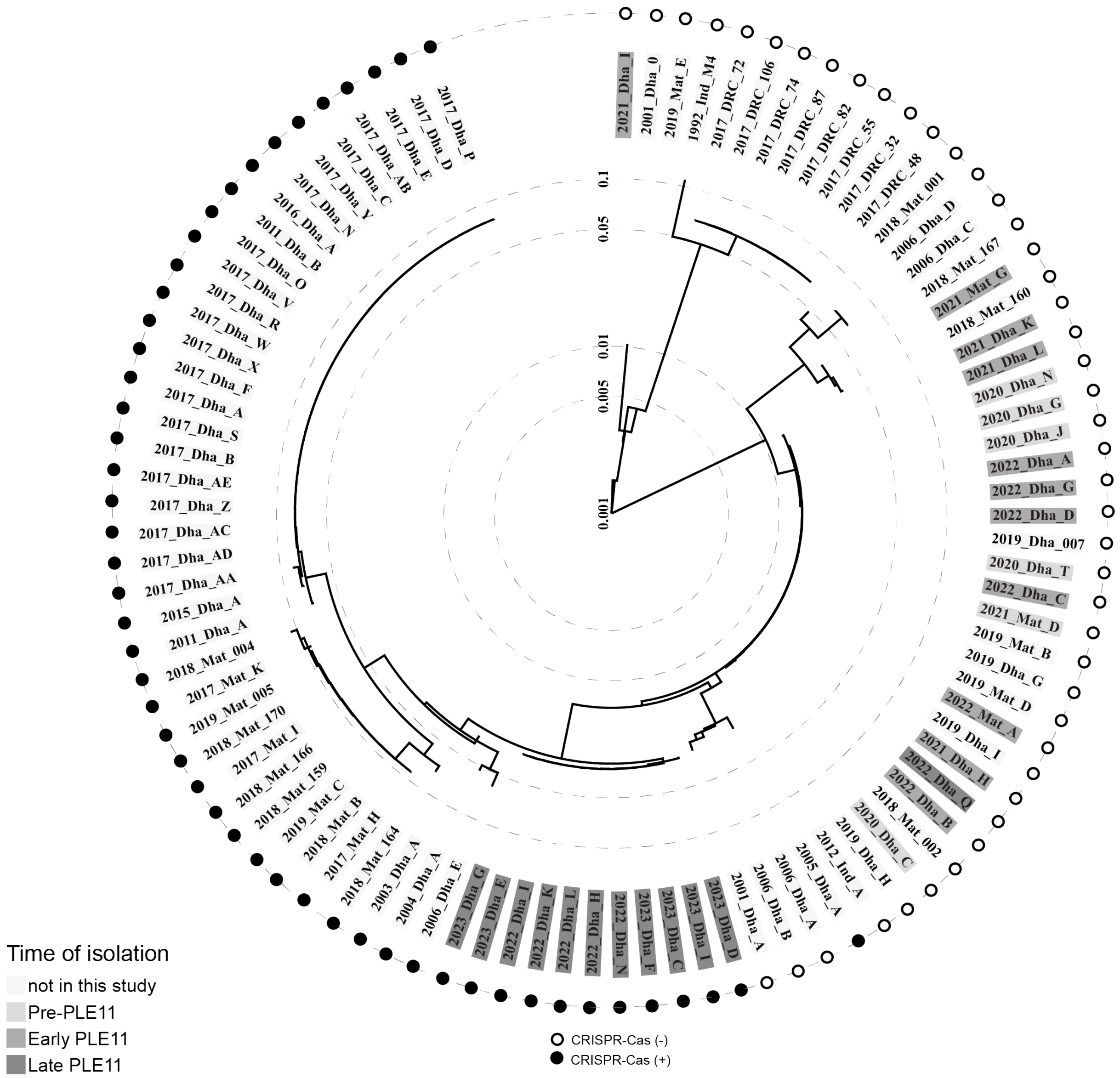


**Extended data Fig. 4: Phylogeny of all sequenced ICP1 isolates**

Phylogeny of 96 whole genome sequenced ICP1 phage isolates (29 from this study and 67 from previously published works (Boyd et al. 2021) based on all translated open reading frames to determine overall similarity (using VipTree's tBLASTx-based algorithm). Filled and empty circles indicate the presence and absence of CRISPR-Cas, respectively. The light gray shading indicates the period of isolation, as the legend indicates. Standardized ICP1 names are shown (including the year and location of isolation: Dhaka Bangladesh, Mat = Mathbaria Bangladesh, Ind = India, DRC = Democratic Republic of Congo).

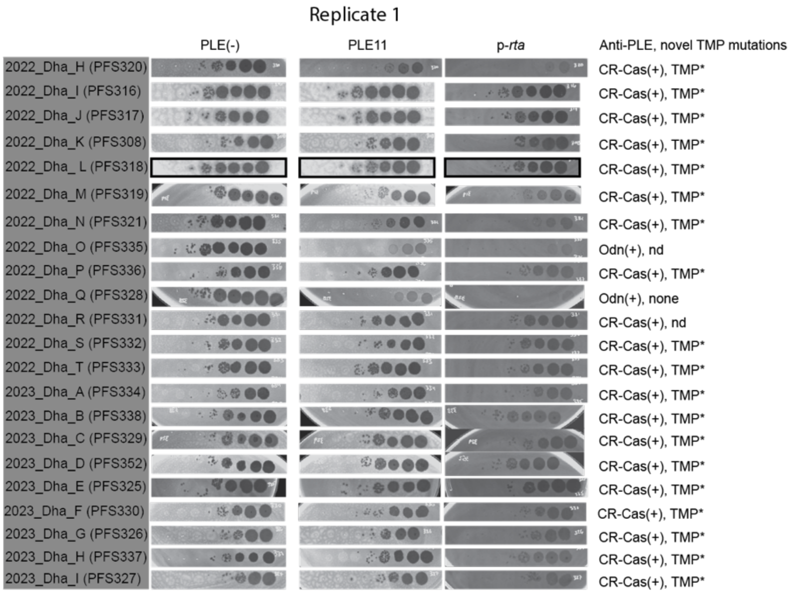


**Extended data Fig. 5. ICP1 isolates from the late-PLE11 period have evolved to overcome PLE11-mediated defense and Rta**

Plaquing of tenfold serially diluted ICP1 phage isolates from the late PLE11 period (July 2022 to December 2023) on lawns of *V. cholerae* strain E7946, E7946 PLE11(+) and E7946 with a low-copy plasmid expressing PLE11 *rta*. The gray background is the bacterial lawn, and the dark spots are zones of killing. Images that were used in the main text are indicated with black boxes. Standardized ICP1 names are shown (including the year and location of isolation: Dha = Dhaka, Mat = Mathbaria, and lab designations (PFS#)). The dark gray shading indicates the period after PLE11 (late-PLE11 period). Anti-PLE mechanisms and TMP mutations are indicated; TMP* refers to either substitution combination of L362P $\pm$ N355S; isolates for which the TMP was not analyzed are noted with “nd” for not determined. Additional biological replicates are included in Supplementary Fig 2.

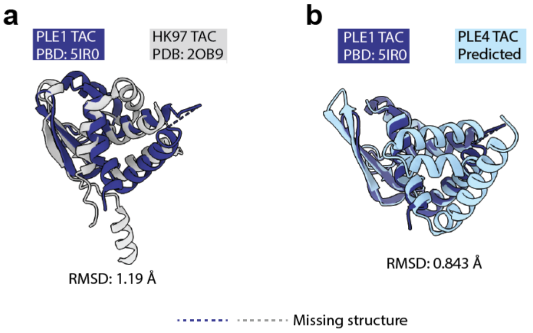


**Extended Data Fig. 6. PLEs encode a tail assembly chaperone structurally similar to a characterized tail assembly chaperone in phage HK97**
**a)** Smith-Waterman alignment between partially solved X-ray diffraction structures of the TAC from PLE1 (PDB:51R0) and HK97 TAC (PDB: 2OB9), a characterized phage TAC.
**b)** Smith-Waterman alignment between partially solved X-ray diffraction structures of the TAC from PLE1 (PDB:51R0) and the predicted structure of the TAC from PLE4. The dotted lines (---) indicate the missing residues.

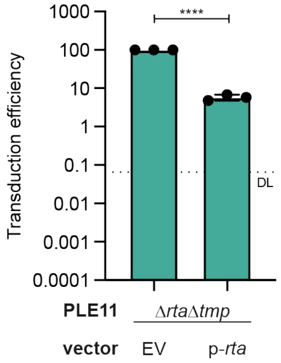


**Extended Data Fig. 7. Expression of Rta *in trans* reduces transduction of PLE11 lacking its own TMP.** Transduction efficiency of Δ*rta*Δ*tmp* PLE11 with an empty (EV) or a plasmid expressing Rta relative to the EV control upon infection by ΔCRISPR-Cas ICP1. The bar represents the mean; each dot represents a biological replicate; the error bars indicate the standard deviation, and DL indicates the detection limit.

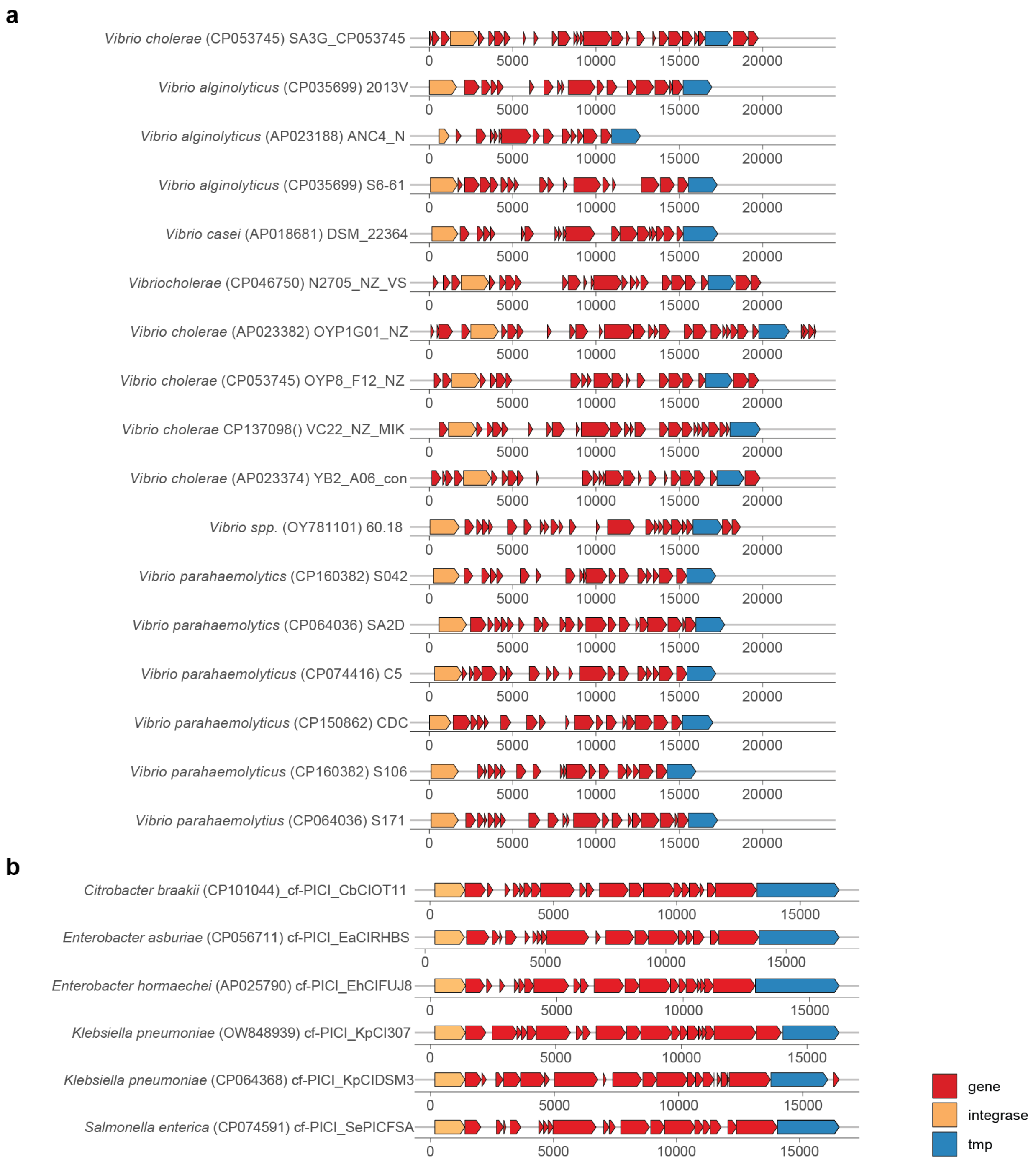


**Extended Data Fig. 8. Putative *tmp* genes are encoded in diverse putative phage satellites.**
**a)** Gene maps of PLE-like elements (identified in LeGault et al. 2022) These elements were queried for homologs of PLE11's tape measure protein (TMP) (using tBLASTn).
**b)** Gene maps of a sampling of cf-PICIs (capsid-forming phage-inducible chromosomal islands, Alqurainy et al. 2023; a family of phage satellites unrelated to PLEs) encoding putative *tmp* genes.
For a and b: Accession numbers are shown within the parentheses. Integrases and TMPs were predicted by homology to known or putative integrases and TMPs using HHPred. All other genes are indicated as arrows for simplicity.
